# EcDNA-borne PVT1 fusion stabilizes oncogenic mRNAs

**DOI:** 10.1101/2025.04.01.646515

**Authors:** Hyerim Yi, Shu Zhang, Jason Swinderman, Yanbo Wang, Vishnupriya Kanakaveti, King L. Hung, Ivy Tsz-Lo Wong, Suhas Srinivasan, Ellis J. Curtis, Aarohi Bhargava-Shah, Rui Li, Matthew G. Jones, Jens Luebeck, Yanding Zhao, Julia A. Belk, Katerina Kraft, Quanming Shi, Xiaowei Yan, Simon K. Pritchard, Frances M. Liang, Dean W. Felsher, Luke A. Gilbert, Vineet Bafna, Paul S. Mischel, Howard Y. Chang

**Author notes:** These authors contributed equally (Hyerim Yi, Shu Zhang). Co-corresponding authors: P.S.M. and H.Y.C.

## Abstract

Extrachromosomal DNA (ecDNA) amplifications are prevalent drivers of human cancers. We show that ecDNAs exhibit elevated structural variants leading to gene fusions that produce oncogene fusion transcripts. The long noncoding RNA (lncRNA) gene *PVT1* is the most recurrent structural variant across cancer genomes, with *PVT1-MYC* fusions arising most frequently on ecDNA. *PVT1* exon 1 is the predominant 5’ partner fused to *MYC* or other oncogenes on the 3’ end. Mechanistic studies demonstrate that *PVT1* exon 1 confers enhanced RNA stability for fusion transcripts, which requires *PVT1* exon 1 interaction with SRSF1 protein. Genetic rescue of MYC-addicted cancer models and isoform-specific single-cell RNA sequencing of tumors reveal that *PVT1-MYC* better supports *MYC* dependency and better activates MYC target genes *in vivo*. Thus, the mutagenic landscape of ecDNA contributes to genome instability and generates chimeric fusions of lncRNA and mRNA genes, selecting *PVT1* 5’ region as a stabilizer of oncogene mRNAs.

## Introduction

Extrachromosomal DNAs (ecDNA) are megabase-sized circular episomes encoding oncogenes and additional genetic elements favoring tumor fitness^1–4^. EcDNA-derived amplifications are detected in 17.1% of tumors from a wide variety of cancer types in adults, with increasing frequency in metastatic tumors^5–7^. EcDNA plays a pivotal role in tumor development and therapy resistance by driving oncogene amplification and tumor evolution^3,5,7–11^. One key feature of ecDNA biology is its proclivity for genomic rearrangements—leading to gene translocations, inversions, insertions and deletions of the ecDNA-borne sequence—which reshapes the tumor genome and contributes to therapeutic resistance and poor prognosis^5,8,12–18^. Recent studies demonstrated that tumors with more advanced stages exhibit increased copy number and higher structural complexity of ecDNA^6,11^. Furthermore, structural variant (SV) burden is particularly elevated within the oncogene loci amplified on ecDNA in the high-risk primary and metastatic breast tumors^19^, underscoring the significant contribution of ecDNA-driven genome instability to aggressive cancer behavior and progression.

Genomic rearrangement can lead to gene fusions when segments of two distinct genes become abnormally joined^20–23^. Gene fusions are a hallmark of many cancers but are rare in normal tissues, making them valuable biomarkers for cancer detection and promising targets for mRNA cancer vaccines^20–28^. Genomic rearrangement with a high SV burden facilitates gene fusions, leading to promoter hijacking or generating fusion transcripts, which disrupts gene regulation and contributes to tumorigenesis and cancer progression^20,23,29–32^. Intriguingly, multiple studies demonstrated a strong association between gene fusions and genomic amplifications across various cancer types^33–36^. Furthermore, a recent finding suggests a potential link between ecDNA and gene fusions, as highly recurrent truncated genes have been identified on ecDNA^37^. Given the growing recognition of ecDNAs as hotspots of genomic rearrangement, understanding their contribution to gene fusions can provide critical insights into cancer biology and new therapeutic strategies.

We hypothesized that ecDNAs may be particularly fertile ground for generating gene fusions that produce gain-of-function transcripts. Unlike chromosomal DNA, ecDNAs are acentric molecules that are asymmetrically inherited during mitosis^13,38^, promoting intratumoral genetic heterogeneity and accelerated evolution. ecDNAs are also a source for ongoing mutagenesis and SVs^7,17,19,39^. This raises the possibility that gene fusions on ecDNA may be more likely to generate functional transcripts with oncogenic potential, which could be rapidly selected in tumors. Oncogene fusion transcripts may serve as functional driver of tumorigenesis^23,29–31^, biomarkers for cancer classification^20,23,32,40^, and RNA templates for cancer vaccine development^41–43^. Despite their potential significance, a systematic and comprehensive understanding of fusion transcripts arising from ecDNA—and their functional impact—remains lacking. Addressing this gap is critical for uncovering new therapeutic targets and advancing cancer diagnostics.

In this study, we systematically investigate the association between gene fusions and amplicon types by integrating whole genome sequencing (WGS) and transcriptome data (RNA-seq) from The Cancer Genome Atlas (TCGA) and Cancer Cell Line Encyclopedia (CCLE) databases, uncovering a strong association between ecDNA and oncogene fusions. We characterize *Plasmacytoma Variant Translocation 1* (*PVT1*) as the most prevalent fusion oncogene arising from ecDNAs across genome and cancer types. Furthermore, we uncover the mechanistic basis for somatic selection of *PVT1* exon 1 as the 5’ gene fusion partner via enhancement of fusion mRNA stability and oncogene function. Together, this study reveals a previously unrecognized connection between ecDNA and oncogene fusion, offering mechanistic insights into the prevalence and oncogenic significance of *PVT1* fusion in cancer.

## Results

### Landscape of gene fusions on ecDNAs

To investigate the association between gene fusions and different amplicon types, we integrated Amplicon Architect (AA) data from whole genome sequencing (WGS) and RNA fusion profiling data from RNA-seq^27^ for 1,825 matched cancer samples across 83 cancer types from TCGA and CCLE databases. This approach enabled a systematic and comprehensive annotation of amplicon types to the breakpoints of 525,648 SVs and 23,546 RNA fusions (**Figure 1A and Table S1**). Fusion transcripts serve as direct evidence of gene fusion, and previous study indicates that 82% of detected fusion transcripts are associated with genomic rearrangements in cancer^20^. To better understand whether genomic rearrangements on ecDNA are a major source of gene fusion transcriptions across cancer, we first analyzed the frequency of co-occurrence of SV (SV burden^19^) and RNA fusion breakpoints (RNA fusion burden) in 100 kb genomic windows across the genome. This analysis revealed a strong alignment between RNA fusion burden peaks and matching SV burden peaks across the genome (**Figure 1B**). We then examined the frequency of this concordance on ecDNA, relative to chromosomal loci, demonstrating a stronger correlation between SV and RNA fusion burden (R=0.7) on ecDNA, than on other focal amplicon types (R≤0.56) or non-focal amplification (No-fSCNA^5^) (R=0.08) (**Figure S1A**). We found that ecDNA-borne RNA fusions account for 14.5% (264/1,825) of all cancers. Among ecDNA(+) cancers (26.2%, 479/1,825), 55.1% (264/479) show evidence of ecDNA-borne RNA fusions, underscoring the strong link between ecDNAs and the production of fusion transcripts.

**Figure 1.**
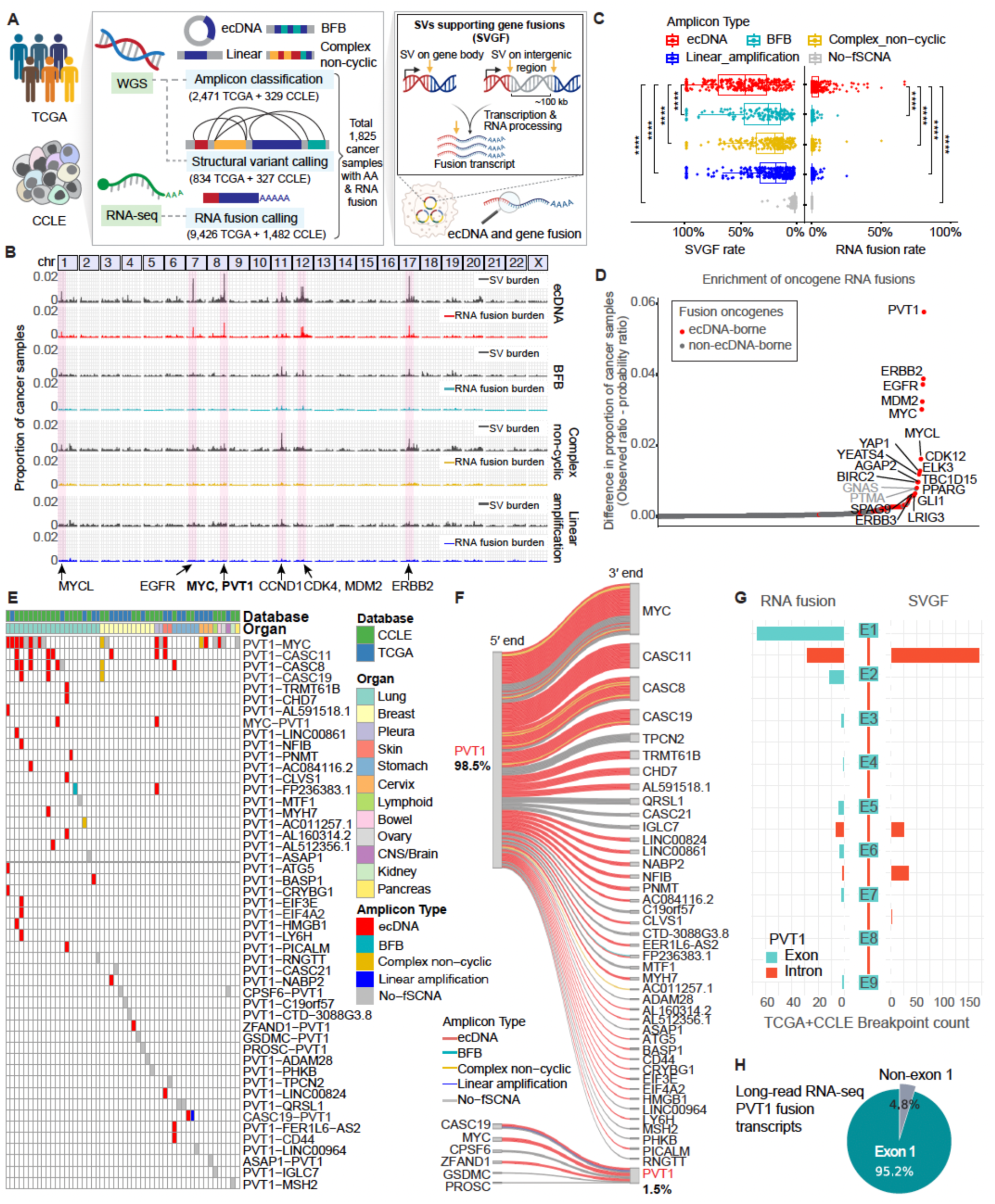
*PVT1* is the most prevalent oncogene fusion enriched by ecDNA. (A) Schematic of the analysis workflow for detecting SV breakpoints supporting gene fusions (SVGF) from whole-genome sequencing (WGS) and RNA fusion breakpoints from RNA-seq, with amplicon type annotation in TCGA and CCLE databases. ecDNA; Extrachromosomal DNA, BFB; breakage-fusion-bridge, No-fSCNA; no focal somatic CN amplification (B) Genome-wide distribution of SV and RNA fusion burden by amplicon type (TCGA and CCLE). SV and RNA fusion burden are shown as the proportion of cancer samples harboring SV or RNA fusion breakpoints within 100 kb genomic windows from each amplicon type, relative to the total number of samples. (C) SVGF rate (left) and RNA fusion rate (right) across different amplicon types (TCGA and CCLE). Rates are defined as the proportion of genes with either SVGF or RNA fusions, relative to the total number of genes within each amplicon interval. Each dot represents a cancer sample classified by amplicon type (****p < 0.0001; two-tailed unpaired t-test). (D) Enrichment of oncogene RNA fusions, ranked by the difference between the observed and expected proportion of cancer samples with either ecDNA-borne RNA fusions (red) or non-ecDNA-borne RNA fusions (gray) (TCGA and CCLE). Each dot represents a fusion oncogene. (E) Distribution of *PVT1* fusion transcripts across cancer types, annotated by amplicon type (TCGA and CCLE). Each row represents a *PVT1* fusion transcript pair, and each column is a cancer sample, clustered by database source and cancer type (organ). Color-coded bars within the heatmap indicate the amplicon type. (F) Position of *PVT1* (5’ or 3’ end) in *PVT1* fusion transcripts (TCGA and CCLE). Each line represents a distinct fusion transcript with unique RNA breakpoints, color-coded by the amplicon type. (G) Distribution of RNA fusion breakpoints (left) and SVGF breakpoints (right) for *PVT1* fusions across exonic and intronic regions of *PVT1* (TCGA and CCLE). (H) Proportion of *PVT1* fusion transcripts containing *PVT1* exon 1, measured by long-read RNA sequencing in ecDNA(+) cell line models and their isogenic pairs where *PVT1* fusion transcripts are detected (COLO320DM, COLO320HSR, PC3DM, SNU16, GBM39KT, and GBM39HSR).

Fusion transcripts can arise from SVs occurring either within a gene body or in intergenic regions, where SVs reposition two genes adjacent to each other, leading to fusion after transcription and RNA processing^23^. To systematically assess the relationship between gene fusion rates and amplicon types, we first defined SVs that contribute to fusion transcript formation as SVGF (SVs supporting gene fusions) based on their breakpoint locations–either within a gene body or in the intergenic region near a gene body (~100 kb)^44–46^ (**Figure 1A**). We then calculated the SVGF rate, defined as the proportion of genes with SVGF relative to the total number of genes within each amplicon interval. EcDNA exhibited the highest SVGF rate compared to other focal and non-focal amplification (No-fSCNA) (**Figure 1C**, left), suggesting that ecDNA are more prone to SVs generating gene fusions. Supporting this idea, RNA-seq analysis revealed that ecDNA also exhibited the highest RNA fusion frequency, calculated as the proportion of genes with RNA fusion breakpoints relative to the total number of genes within each amplicon interval, compared to cancers with other mechanisms of gene amplification (**Figure 1C**, right). Using long-read DNA and RNA sequencing in multiple ecDNA(+) cell line models, we further validated the precise SV breakpoints on ecDNA encoding highly expressed fusion transcripts across different cell lines (**Figure S1B and Table S1**). Notably, the top expressed fusion mRNA transcripts in each cell line were nearly always from gene fusions on ecDNA. In line with this, the overall expression levels of ecDNA-borne fusion transcripts were elevated compared to non-ecDNA-borne fusion transcripts (**Figure S1C**), reinforcing ecDNA as a hotspot for SV-driven gene fusions and consequent amplification of fusion transcript expression.

EcDNAs are more prone to harbor activating mutations that confer a selective advantage to cancer cells, facilitating positive selection^38,47,48^. To identify specific gene fusions strongly associated with ecDNA, we checked the genes with SV and RNA fusion burden peaks across genome (**Figure 1B**). In ecDNA(+) cancers, both SV and RNA fusion burden peaked at well-known ecDNA-amplified oncogenes such as *EGFR, MYC/PVT1, MDM2,* and *ERBB2* (**Figure 1B**). Our analysis of recurrent oncogene RNA fusions revealed that the most frequently observed oncogene RNA fusions in ecDNA-borne RNA fusions were predominantly ecDNA-amplified oncogenes^7,37^ (**Figure S1D**). In contrast, non-ecDNA-borne RNA fusions were enriched in a distinct set of oncogenes not commonly amplified on ecDNA (**Figure S1D**). To further test whether ecDNA-borne RNA fusions preferentially enrich oncogene fusions compared to non-ecDNA-borne RNA fusions, we calculated the difference between observed and expected probability ratios of cancer samples with each oncogene RNA fusion in ecDNA-borne vs non-ecDNA-borne RNA fusion samples. The expected probability ratio was derived by assuming a random distribution of RNA fusions, weighted by the proportion of ecDNA-borne vs non-ecDNA-borne RNA fusions relative to the total RNA fusions. Strikingly, ecDNA-borne RNA fusion samples showed a strong preferential enrichment of oncogene RNA fusions (**Figure 1D**, red dots), Collectively, our data demonstrate that structural rearrangements on ecDNA frequently give rise to oncogene fusions, leading to their elevated expression and selective enrichment, highlighting ecDNA’s role as a key driver of oncogene fusion transcript formation in cancer.

### EcDNA-borne *PVT1-MYC* fusion mRNA

The *MYC/PVT1* locus exhibited the highest frequency of both SV and RNA fusion burden across the entire cancer genome (**Figures 1B and S1E**). *PVT1* suffers frequent gene fusion involving multiple fusion partners across human cancers^20,49,50^. *PVT1* is located 50 kb 3’ of the *MYC* oncogene on human chromosome 8q24^51,52^. *PVT1* encodes a long non-coding RNA (lncRNA), which is expressed at low levels in normal tissues, but its upregulation is strongly associated with tumor onset, progression, and poor prognosis^49,50,53,54^. Multiple studies have linked *PVT1* fusions to 8q24 copy-number amplification^4,50,55,56^. For instance, *PVT1-MYC* fusion is detected in 60% of *MYC*-amplified group 3 medulloblastomas, characterized by *MYC* ecDNA amplification^56–58^. However, a systematic association between *PVT1* fusions and ecDNA has not been previously addressed.

Our analysis unveils that *PVT1* gene fusion was selectively enriched in ecDNA(+) cancers, while peaks at some other ecDNA-associated oncogenes, such as *CCND1* and *ERBB2*, were distributed across multiple amplicon types (**Figure 1B**). *PVT1* fusion is enriched in cancers with ecDNA-borne RNA fusions by 4.4-fold compared to non-ecDNA-borne RNA fusion cancers (**Figure S1D**). Furthermore, *PVT1* is the top recurrent oncogene RNA fusion in ecDNA(+) cancers (**Figures 1B and 1D**) and *PVT1* exhibits the highest number of fusion transcript among oncogenes across all cancer samples, with ecDNA accounting for 72.3% of all detected *PVT1* fusion transcripts (**Figure S1F**). Moreover, when *PVT1* is amplified on ecDNA, it generates the higher number of fusion variants, and overall, genes on ecDNA tend to produce more diverse fusion species than those on other amplicon types (**Figure S1G**), highlighting the role of ecDNA in generating oncogene RNA fusions.

Next, we investigated the distribution of *PVT1* fusion transcripts across cancer types and their association with amplicon types. *PVT1* fusions were detected broadly across cancers, with the highest prevalence in lung cancers (7%; 20/287), where most *PVT1* fusions are ecDNA-associated (65%; 13/20) (**Figure 1E**). In contrast, breast cancers showed lower *PVT1* fusion association with ecDNA and higher association with No-fSCNA, suggesting tissue-specific patterns in *PVT1* fusion formation (**Figure 1E**). Notably, ecDNA-amplified genes such as *MYC, CASC11*, and *CASC19*, which frequently undergo RNA fusions and co-amplified with *PVT1* are *PVT1* fusion partners (**Figure 1E and Table S1**), reinforcing the role of ecDNA as a platform that promotes prevalent oncogenic RNA fusions. Furthermore, expression levels of *PVT1* fusion transcripts were significantly higher in cancers with ecDNA-borne RNA fusion compared to non-ecDNA-borne RNA fusion cancers (**Figure S1H**), highlighting the potential of *PVT1* fusion transcripts as biomarkers for ecDNA(+) cancers. Indeed, by combining *PVT1* fusion transcript expression with *PVT1* copy number gain (CN > 10), ecDNA detection accuracy was improved to 95%, whereas using either metric alone yielded lower accuracies of 74.5% and 82.1%, respectively (**Figure S1I**).

Our integrated analysis of TCGA and CCLE showed that 98.5% of *PVT1* fusion transcripts have *PVT1* fusion located at their 5’ end (**Figure 1F**), consistent with prior anecdotal reports^50^. These recurrent SV and RNA fusion breakpoints are primarily located in *PVT1* exon 1 and intron 1, generating mature fusion transcripts that retain exon 1 of lncRNA *PVT1* spliced onto multiple candidate 3’ coding sequences (**Figure 1G**). Leveraging long-read DNA and RNA-sequencing in ecDNA(+) cell line models, we precisely mapped the *PVT1* fusion breakpoints that most frequently located in exon 1 and intron 1 leading to 95.2% of *PVT1* fusion transcripts harboring exon 1 (**Figures 1H and S1J**). Considering that *PVT1* exon 1 comprise 7–12% of the *PVT1* lncRNA isoforms but are observed in 95% of the amplified fusion transcript, this result represents an 8–14-fold enrichment to retain *PVT1* exon 1. Such recurring mutational “hotspot” is indicative of gain of function for an oncogene product^59^, reflecting positive selection for *PVT1* exon 1 in tumor evolution facilitated by the asymmetric inheritance of ecDNA. Together, these results suggest that ecDNA-mediated genomic rearrangements drive 5’ end *PVT1* exon1 fusions, making them a predominant form of oncogenic fusion events in cancer.

### *PVT1* exon 1 fusion enhances mRNA stability

To investigate the molecular function of *PVT1* exon 1 fusion, we utilized a cell line model carrying *PVT1-MYC* fusion generated by ecDNA, which is the most prevalent form of *PVT1* fusion across cancer (38.5%; 20/52) (**Figure 2A**). In the colorectal cancer cell line COLO320DM, ecDNAs contain multiple copies of the *PVT1-MYC* fusion, where the *MYC* promoter and exon 1 are replaced by the *PVT1* promoter and exon 1 (**Figure 2A**). This rearrangement produces a mature *PVT1-MYC* fusion RNA, where *PVT1* exon 1 is fused to *MYC* exon 2–3, replacing the 5’ UTR of canonical *MYC*^4^. Intriguingly, in its isogenic paired cell line, COLO320HSR, *MYC* is amplified on chromosome not ecDNA, without either the amplification or fusion of *PVT1* 5’ region^4^. Consistent with this, we demonstrated by metaphase DNA FISH that the majority of *PVT1* exon 1 and *MYC* signal colocalize on ecDNA in COLO320DM while we rarely observed colocalized signal on homogenous staining regions (HSR) chromosome in COLO320HSR (**Figures 2B and S2A**). While some colocalization occurs at rare linear integration sites, it is absent in canonical HSR regions, resulting in *PVT1-MYC* fusion being largely absent in COLO320HSR (**Figure S2A**).

**Figure 2.**
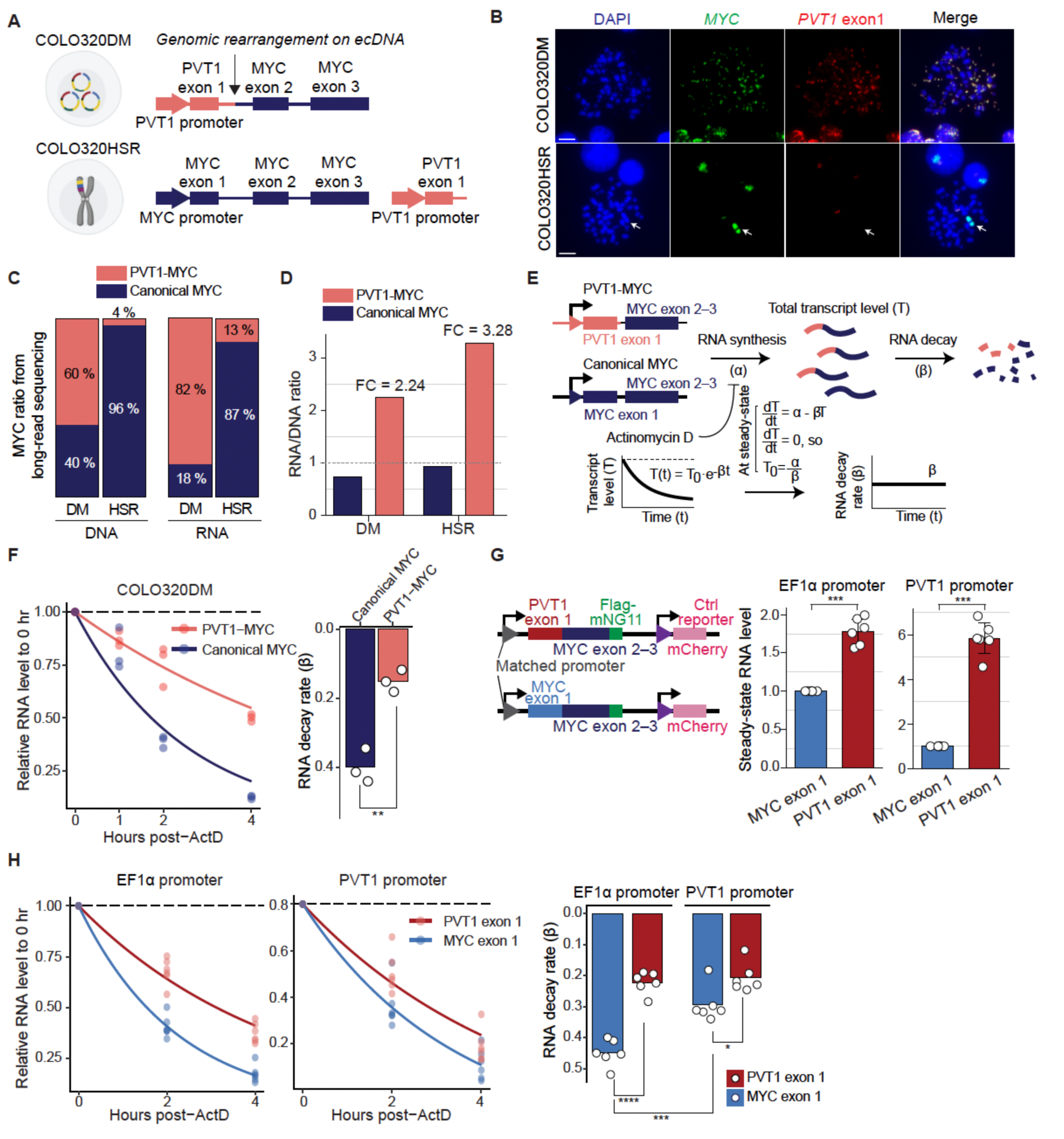
*PVT1* exon 1 fusion enhances oncogene mRNA stability. (A) Schematic of the genomic rearrangement on ecDNA in COLO320DM generating *PVT1-MYC* fusion and the restored canonical *MYC* sequence in the isogenic cell line, COLO320HSR. (B) Representative metaphase DNA FISH images of COLO320DM and COLO320HSR cell lines with FISH probes targeting *MYC* and *PVT1* exon 1 region. Scale bars, 10 μm. (C) Proportion of DNA copy number (left) and RNA abundance (right) of *PVT1-MYC* and canonical *MYC* in COLO320DM and COLO320HSR cell lines, measured by long-read sequencing. (D) Normalized RNA abundance relative to DNA copy number of *PVT1-MYC* and canonical *MYC* transcripts in COLO320DM and COLO320HSR cell lines. Fold change (FC) between *PVT1-MYC* and canonical *MYC* in each cell line is indicated above the *PVT1-MYC* bars. (E) Schematic of RNA stability assay and a mathematical modeling for calculating RNA decay rate (β), based on the decrease in steady-state transcript levels over time following transcription inhibition. (F) RNA stability measurement and decay rate analysis of endogenous *PVT1-MYC* and canonical *MYC* transcripts in COLO320DM cells using RT-qPCR (n=3). Cells were harvested at 0, 1, 2, and 4 hr after actinomycin D treatment (4 ng/mL). Left: Each dot represents relative RNA abundance normalized to 0 hr and *GAPDH* internal control RNA, with fitted lines based on decay rates from mathematical modeling. Right: RNA decay rate of canonical *MYC* and *PVT1-MYC*. Each dot represents a decay rate for each replicate (**p < 0.002; two-tailed t-test). (G) Left: Schematic of the reporter assay system with either *PVT1* exon 1 or *MYC* exon 1 fusion. Right: Steady-state RNA levels of MYC-Flag-mNG11 reporter transcripts driven by either the *EF1α* or *PVT1* promoter, normalized to mCherry internal control RNA (n=6). RNA levels were measured by RT-qPCR 48 hr post-transfection with equal amounts of plasmid DNA in COLO320DM cells (***p < 0.0002; two-tailed t-test). (H) RNA stability measurement of reporter transcripts shown in Figure 2G. COLO320DM cells that were transfected with the equal amounts of plasmid DNA, were harvested at 0, 2, and 4 hr after actinomycin D treatment (4 ng/mL) 48 hr post-transfection (n=6). Left: Each dot represents relative RNA abundance normalized to 0 hr and *GAPDH* internal control, with fitted lines based on decay rates from mathematical modeling. Right: RNA decay rates of reporter RNA stability assay. Each dot represents a decay rate for each replicate (*p < 0.03, ***p < 0.0002, ****p < 0.0001; two-tailed t-test).

To investigate the impact of *PVT1-MYC* fusion on gene expression, we compared DNA copy number and RNA expression levels of *PVT1-MYC* and canonical *MYC* using long-read single molecule sequencing (**Figure S2B**), precisely identifying and quantifying mRNA isoforms. In COLO320DM, *PVT1-MYC* mRNA accounts for 82% of total *MYC* transcripts, whereas in COLO320HSR, canonical *MYC* is the major *MYC* isoform at 87% of total *MYC* mRNA (**Figure 2C**). These findings support the preferential association of *PVT1-MYC* fusion with ecDNA (**Figures 1E–F**). In line with this idea, our analysis in TCGA and CCLE databases revealed that *PVT1-MYC* RNA fusion was 6-fold enriched in cancer samples with ecDNA-borne RNA fusions (3.9%; 10/264) compared to its frequency in cancers with non-ecDNA-borne RNA fusion (0.6%; 10/1561). Notably, after normalizing for DNA copy number, steady state RNA level of *PVT1-MYC* mRNA was ~2–3-fold higher than that of canonical *MYC* mRNA in both COLO320DM and COLO320HSR cells (**Figure 2D**), suggesting that *PVT1-MYC* mRNA expression is elevated beyond DNA copy number.

To test whether *PVT1* exon 1 fusion increases mRNA transcription or stability of its fusion partner, we developed luciferase reporter constructs (nLuc) with or without PVT1 exon 1 under the matched promoter to control for transcription efficiency (**Figure S2C**). Steady-state reporter RNA levels were assessed by normalizing to an internal control reporter RNA (fLuc), which was expressed under a separate promoter on the same plasmid, to account for transfection efficiency (**Figure S2C**). *PVT1* exon 1 fusion resulted in a 2–3-fold increase in reporter mRNA levels compared to the non-fused reporter under the same promoter (minimal promoter^4^) (**Figure S2C**, left). This fold change closely matched the difference in endogenous RNA expression between *PVT1-MYC* vs canonical *MYC* mRNAs (**Figure 2D**), suggesting that post-transcriptional mechanisms contribute to the elevated RNA level of *PVT1-MYC*. Furthermore, when *PVT1* exon 1 fusion was combined with the *PVT1* promoter–the predominant *PVT1* fusion pattern across cancer (**Figures 1G and S1J**)–reporter RNA level increased more than 10-fold (**Figure S2C**, right). This suggests a synergistic effect between transcriptional and post-transcriptional regulation, reinforcing the functional advantage of *PVT1* 5’ end fusion in cancer.

To address whether *PVT1* exon 1 fusion regulates RNA stability, we measured RNA half-life of endogenous *PVT1-MYC* and canonical *MYC* mRNAs by blocking transcription using actinomycin D and tracking total RNA abundance over time. RNA decay rates were then estimated based on the observed RNA abundance changes using mathematical modeling (**Figure 2E and Methods**). *PVT1-MYC* exhibited higher RNA stability compared to canonical *MYC* in both COLO320DM and COLO320HSR cells (**Figures 2F and S2D**), suggesting that the sequence-mediated regulatory mechanisms contribute to its increased stability. Additionally, the 2–3-fold difference in RNA decay rates between *PVT1-MYC* and canonical *MYC* can nearly fully explain the comparable fold difference in steady state endogenous RNA levels measured by long-read sequencing (**Figures 2D and S2E**), further supporting RNA stability as the major factor underlying the elevated expression of *PVT1-MYC*.

To further investigate whether *PVT1* exon 1 fusion confers RNA stability to *MYC* coding sequence, we generated reporter constructs that fused either with *PVT1* exon 1 or *MYC* exon 1 to *MYC* exon2–3-Flag-mNG11 reporter, mimicking the endogenous condition (**Figure 2G**). Consistent with our previous luciferase-based reporter experiments (**Figure S2C**), the *PVT1* exon 1-fused reporter exhibited higher steady-state RNA expression than the *MYC* exon 1-fused reporter, and the combined fusion of *PVT1* promoter and exon 1 led to an even greater increase in RNA expression (**Figure 2G**). Under the control of transcription efficiency using matched promoters, *PVT1* exon 1-fused reporter exhibited higher RNA stability compared to *MYC* exon 1-fused reporter, demonstrating that *PVT1* exon 1 fusion confers RNA stability (**Figure 2H**). The fold change in RNA decay rates between the *PVT1* exon 1-vs *MYC* exon 1-fused reporters under the *EF1α* promoter closely matched the fold difference in their observed steady-state RNA levels (**Figures 2G and S2F**, left), reinforcing *PVT1* exon 1’s major role in stabilizing RNA and driving increased RNA expression. Interestingly, when transcribed under the *PVT1* promoter, *MYC* exon 1-fused RNA exhibits lower decay rate compared to its expression under the *EF1α* promoter (**Figure 2H**), indicating an additional contribution of *PVT1* promoter to RNA stability.

### RNA-protein interactome reveals SRSF1 binding in *PVT1*-mediated RNA stabilization

To investigate the molecular mechanism underlying *PVT1* exon 1-mediated RNA stabilization, we explored the protein interactome of *PVT1-MYC* and canonical *MYC* RNA using ChIRP-MS (Comprehensive identification of RNA-binding proteins by mass spectrometry), a technique that identifies RNA-associated proteome by crosslinking and pulldown of RNA-protein complexes followed by mass spectrometry^60^ (**Figure 3A**). We leveraged the COLO320DM and COLO320HSR cell lines, where *PVT1-MYC* accounts for 82% of total *MYC* transcripts in COLO320DM, while canonical *MYC* represents 87% of *MYC* transcripts in COLO320HSR (**Figure 2C**). Using probes against *PVT1* exon 1 and exon 1–3 of canonical *MYC* to capture total *MYC* transcript isoforms in each cell line (**Table S2**), we performed a comparative analysis of the *MYC* mRNA-associated proteome between the two cell lines. We identified proteins specifically enriched in COLO320DM MYC ChIRP-MS, despite the comparable protein expression in both cell lines, suggesting preferential binding to *PVT1-MYC* transcripts (**Figures 3B–C**). Specifically, we found multiple members of the SR (Serine/Arginine-Rich Splicing Factor) protein family, which are well-known RNA binding proteins which play key roles in pre-mRNA splicing and RNA metabolism^61–64^ (**Figure 3B and Table S3**). Using publicly available eCLIP (Enhanced crosslinking and immunoprecipitation) data–a high-throughput sequencing method used to identify RNA-binding protein (RBP) interaction sites on RNA transcripts^65^, we found that the SR protein SRSF1, but not SRSF7, strongly occupies *PVT1* exon 1 across two independent cell lines, indicating a reciprocal interaction between SRSF1 and *PVT1* exon 1 (**Figures 3D and S3A**). SRSF1 is known to be overexpressed in cancer and promotes cell transformation^61,66^, and consistent with this, COLO320DM cells displayed higher SRSF1 protein expression compared to non-ecDNA-related colorectal cancer (HCT116) and non-cancer (IMR90) cell lines (**Figure 3C**). Notably, SRSF1 binding to RNA has been reported to enhance RNA stability^62^, making SRSF1 a strong candidate contributing to *PVT1* exon 1-mediated RNA stabilization.

**Figure 3.**
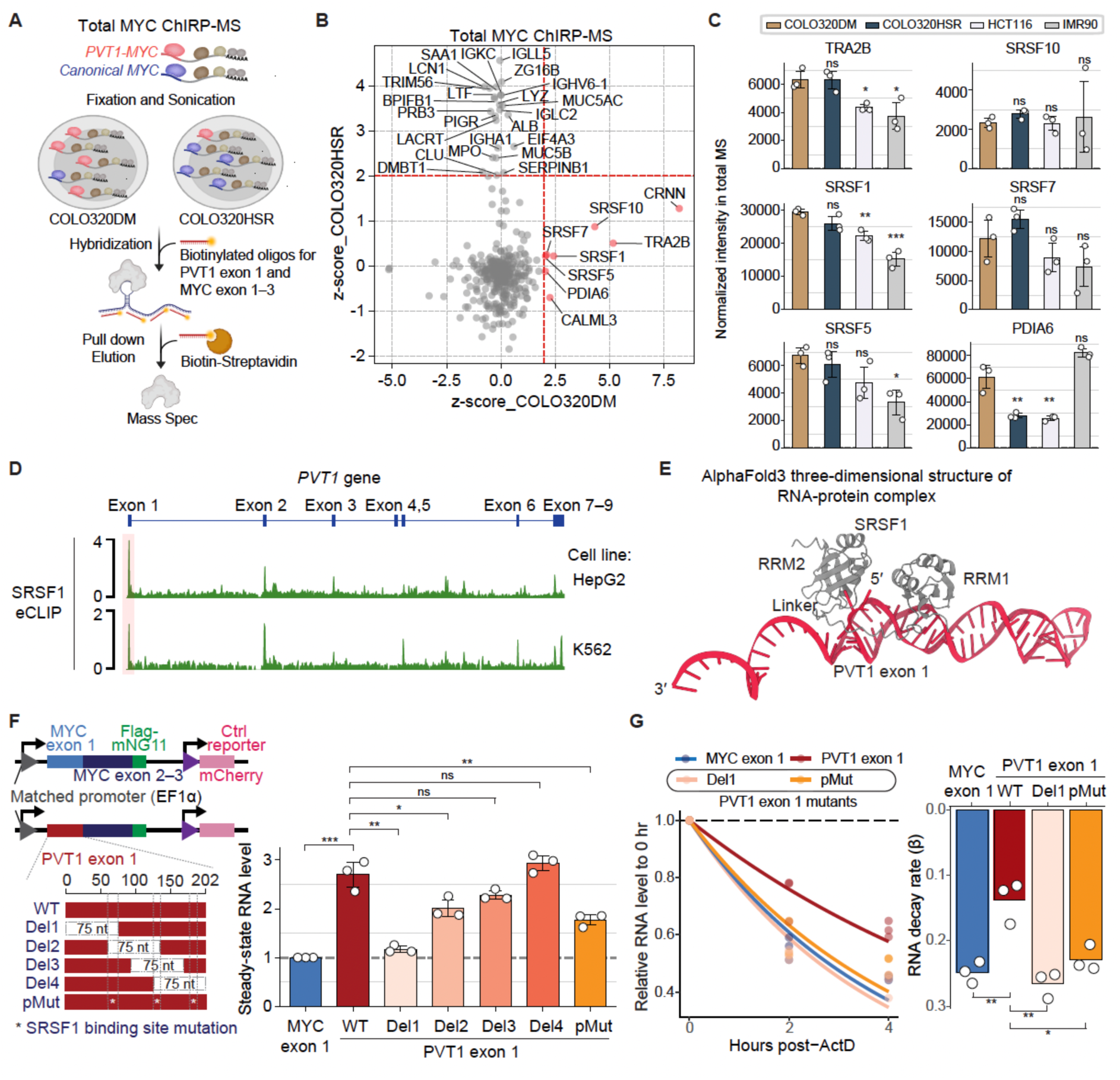
SRSF1 binding sites in PVT1 exon 1 play key roles in RNA stability. (A) Schematic of total *MYC* ChIRP-MS using probes targeting *PVT1* exon 1 and *MYC* exon 1–3 to pulldown total MYC transcripts in COLO320DM and COLO320HSR cell lines. (B) Scatter plot of z-scores comparing total *MYC* ChIRP-MS hits in COLO320DM (x-axis) and COLO320HSR (y-axis) cells. ChIRP-MS hits (z-score > 2) are labeled with gene names (red dashed lines; z-score = 2), with enriched hits in COLO320DM cells highlighted as red dots. (C) Normalized protein abundance from total MS of COLO320DM-enriched *MYC* ChIRP-MS hits (Figure 3B) across COLO320DM, COLO320HSR, HCT116 (non-ecDNA colorectal cancer cell line), and IMR90 (non-cancer cell line) (n=3). P values were calculated by comparing protein expression, relative to COLO320DM (*p < 0.03, **p < 0.002, ***p < 0.0002; two-tailed t-test). (D) Read density tracks of SRSF1 eCLIP across exonic and intronic regions of *PVT1* in HepG2 and K562 cell lines, with an enriched peak at *PVT1* exon 1 highlighted in a pink box. (E) *De novo* AlphaFold3-predicted three-dimensional structure of the interaction between two RRMs of SRSF1 and *PVT1* exon 1 mRNA (62–141 bp). (F) Left: Schematic of the mutant reporter assay system either with *MYC* exon 1 or *PVT1* exon 1 wild type (WT), deletion mutants (Del1–4), or point mutant targeting SRSF1 binding motifs (pMut), driven by the *EF1α* promoter. Right: Steady-state RNA levels of reporter transcripts, normalized to mCherry internal control RNA in HEK293T cells (n=3). RNA levels were measured by RT-qPCR 48 hr post-transfection with equal amounts of plasmid DNA (*p < 0.03, **p < 0.002, ***p < 0.0002; two-tailed t-test). (G) Left: RNA stability measurement of mutant reporter transcripts (Del 1 and pMut) compared to *PVT1* exon 1 WT (positive control) and *MYC* exon 1 (negative control) reporter from Figure 3F. HEK293T cells were transfected with equal amounts of plasmid DNA, and harvested at 0, 2, and 4 hr after actinomycin D treatment (4 ng/mL) 48 hr post-transfection (n=3). Each dot represents relative RNA abundance normalized to 0 hr and *GAPDH* internal control, with fitted lines based on decay rates from mathematical modeling. Right: RNA decay rates of reporter RNA stability assay in HEK293T cells. Each dot represents a decay rate for each replicate (*p < 0.03, **p < 0.002; two-tailed t-test).

### SRSF1 binding sites in *PVT1* exon 1 play key roles in RNA stability

To characterize the RNA-protein interaction between *PVT1* exon 1 and SRSF1, we computationally modeled the three-dimensional structure of the protein-RNA complex of SRSF1 and mRNA of *PVT1* exon 1 using *de novo* structure prediction from AlphaFold 3^67^ and manual refinement^68^ (**Methods**). The structural analysis revealed that the two RNA Recognition Motif (RRM) domains of SRSF1 interact with the 5’ end of *PVT1* exon 1 where the known SRSF1 binding motif (GAGGA) is located, consistent with prior reports indicating that SRSF1 preferentially binds purine (A/G)-rich sequences in exons to promote exon inclusion (**Figures 3E and S3B**).

To assess the functional contribution of SRSF1 binding to *PVT1* exon 1-mediated RNA stabilization, we conducted scanning a mutagenesis experiment to selectively abolish SRSF1 binding to *PVT1* exon 1 of reporter mRNA while avoiding promiscuous effects from globally affecting SRSF1’s role as a splicing factor. Based on the previously reported SRSF1 binding motifs and our AlphaFold3-based predictions, we introduced point mutations in three putative SRSF1 binding sites by replacing GGA with UUA to disrupt SRSF1 binding. The impact of these mutations was examined by DeepCLIP^69^, a deep learning model trained on ENCODE SRSF1 eCLIP data, which showed a decrease in SRSF1 binding scores at the mutated sites (**Figure S3B**). Along with this point mutant reporter (pMut), we generated the deletion mutants by tiling 75-nt segments spanning *PVT1* exon 1, removing sequences that cover the mutation sites to define the minimal sequence element responsible for RNA stabilization (**Figure 3F**).

Using these mutant reporter constructs, we measured steady-state reporter RNA expression in HEK293T cells to investigate whether SRSF1 binding influences PVT1 exon 1-mediated mRNA stabilization and whether this effect is cell type-independent, as SRSF1 binding to *PVT1* exon 1 was commonly observed across different cell lines (**Figure 3D**). We found that deletion of the 5’ end 75 nt of *PVT1* exon 1 (Del1), which contains the AlphaFold3-predicted SRSF1 binding site, completely abolished *PVT1* exon 1-mediated RNA stabilization (**Figures 3F–G**). In line with this result, reporter construct of *PVT1* exon 1 bearing point mutations of the SRSF1 binding sites (pMut) exhibited a significant decrease in reporter mRNA level (**Figure 3F**). Consistently, Del1 and pMut reporters exhibited decreased mRNA stability (**Figure 3G**). Consistent with the idea that an SRSF1 binding sites in the first 193 nucleotides of *PVT1* RNA is important in its oncogenic fusions, RNA breakpoints are absent in this region in *PVT1*-fused transcripts detected in ecDNA(+) cancer cell line models, while RNA breakpoints sharply accumulate distal to 200+ nt of *PVT1* (**Figure S3C**). Despite lncRNA generally being poorly conserved^70,71^, the 5’ end of *PVT1* exon 1 is known to be highly conserved^49,52^, exhibiting a perfect sequence match between mice and humans. This pattern further supports the significance of 5’ end of *PVT1* exon 1 in RNA stabilization, conferring a functional advantage in cancer-associated *PVT1* exon1 fusions.

### *PVT1* fusion enhances the oncogenic function of *MYC*

To investigate how *PVT1-MYC* functions cancer cells, we examined the ability of *PVT1-MYC* to substitute for oncogenic function of *MYC* in *MYC*-addicted cancer cells^72–75^. *MYC*-addicted EC4 cancer cells were engineered to have *MYC* expression under the control of a doxycycline regulatable promoter (Tet-off), enabling depletion of canonical *MYC* upon doxycycline treatment^76^. We transfected equal amounts of DNA constructs encoding either *PVT1-MYC* or canonical *MYC* under the same promoter 24 hr before *MYC* depletion, and measured cell viability 24 hr after endogenous *MYC* depletion in *MYC*-addicted EC4 cancer cells (**Figure 4A**). We found that *PVT1-MYC* more efficiently rescues cancer cell viability in response to *MYC* depletion compared to canonical *MYC* (**Figure 4B**). Replacement of the *EF1α* promoter with the *PVT1* promoter further increased cell viability (**Figure 4B**), consistent with the observation that *PVT1-MYC* showed elevated expression under the *PVT1* promoter (**Figures S2C and 2G**). Furthermore, a 75-nt deletion at the 5’ end (Del1) or point mutations (pMut) of *PVT1* exon 1 that impair SRSF1 binding abrogated the ability of *PVT1-MYC* to rescue *MYC* silencing (**Figure 4B**). These results demonstrate that *PVT1-MYC* can rescue the loss of MYC oncogenic activity but requires the SRSF1 binding site at the 5’ end of *PVT1* exon 1.

**Figure 4.**
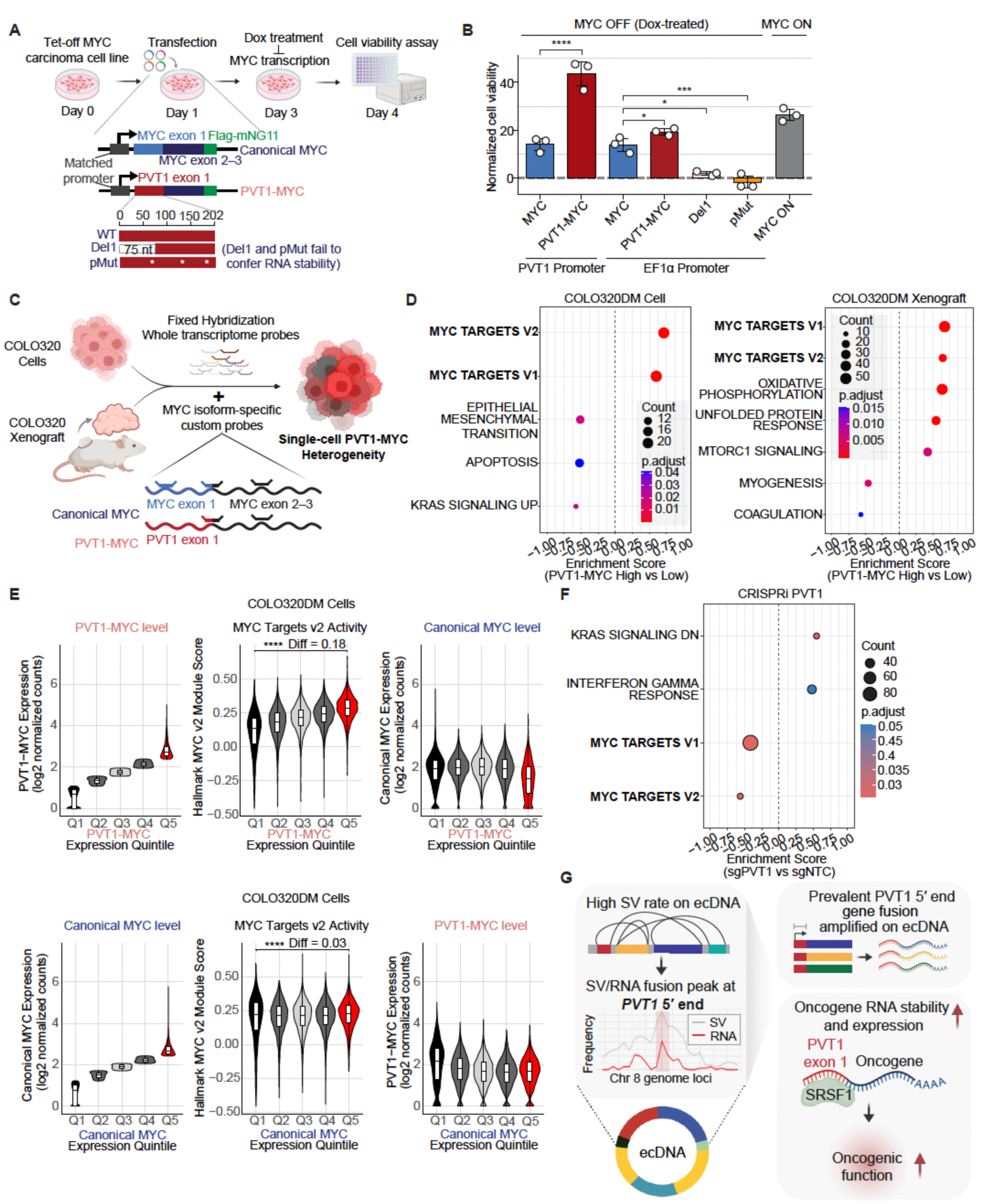
*PVT1* fusion enhances the oncogenic function of *MYC*. (A) Schematic of the *MYC* rescue experiment using the Tet-off *MYC* carcinoma cell line (EC4) and the canonical *MYC*, *PVT1-MYC* WT and mutant reporter constructs (Del 1 and pMut). (B) Normalized cell viability relative to the *GFP* negative control in the *MYC* rescue experiment (n=3) (*p < 0.03, ***p < 0.0002, ****p < 0.0001; one-way ANOVA test). (C) Schematic of Flex scRNA-seq with custom probes targeting *PVT1-MYC* and canonical *MYC* in COLO320DM and COLO320HSR cells and xenograft mice *in vivo* models. (D) Significantly enriched Hallmark pathways in *PVT1-MYC* high vs low cells from Flex scRNA-seq in COLO320DM cell and xenograft models. Pathways are visualized as dot plots indicating gene count and adjusted p-value. (E) Violin plots displaying dose-dependent MYC targets v2 activity and expression level of *MYC* isoforms across expression quintiles of *PVT1-MYC* (top) and canonical *MYC* (bottom) in COLO320DM cells. Difference between mean of the highest quintile (Q5) and lowest quintile (Q1) is presented as Diff (****p < 0.0001; Wilcoxon rank-sum-test). (F) Significantly enriched Hallmark pathways in CRISPRi sgPVT1 vs sgNTC (non-targeting control) RNA-seq in COLO320DM cells. Pathways are visualized as dot plots indicating gene count and adjusted p-value. (G) Schematic model of ecDNA-borne *PVT1* fusion and its oncogenic function.

As the *MYC* oncogene encodes a transcription factor that regulates downstream gene expression, we next investigated whether the *PVT1-MYC* can regulate the expression of MYC target genes. Using Flex single-cell RNA-seq (scRNA-seq)^77,78^, a whole-transcriptome profiling method that allows detection of transcript isoforms in single cells via isoform-specific probes (**Table S2**), we measured *MYC* isoform-specific RNA expression of single COLO320DM and COLO320HSR cancer cells both *in vitro* and *in vivo* xenografts (**Figures 4C and S4A**). Due to lack of centromeres, ecDNAs are randomly inherited during cell division, leading to copy number heterogeneity in cell population^13,38^. COLO320DM cells, which contains ecDNA, show a high level of heterogeneity in *PVT1-MYC* expression among cells, consistent with variable ecDNA copy numbers due to uneven ecDNA segregation^38^. Leveraging the cell-to-cell variation detection for *PVT1-MYC* RNA expression in COLO320DM, we examined the pathways regulated by *PVT1-MYC* expression by binning cells based on its expression.

Cells with high *PVT1-MYC* expression (highest quintile; Q5) showed significantly upregulated MYC target genes in gene pathway enrichment analysis compared to cells with low *PVT1-MYC* expression (lowest quintile; Q1), under both *in vitro* and *in vivo* xenograft settings (**Figures 4D, S4B and Table S4**). Additionally, MYC target scores showed a stronger dose-dependent increase based on *PVT1-MYC* expression compared to canonical *MYC* expression both *in vitro* and *in vivo* (**Figure 4E and S4C**), indicating that *PVT1-MYC* more potently activates MYC target genes. In COLO320HSR cells, where *PVT1-MYC* expression is low, canonical *MYC* expression and MYC target scores displayed less variation than in COLO320DM cells (**Figure S4D**). These results suggests that when highly expressed, *PVT1-MYC* dominates the oncogenic function of *MYC*. Supporting this idea, transcriptional silencing of *PVT1* promoter by CRISPR interference (CRISPRi) in COLO320DM cells significantly decreased MYC target genes in RNA-seq by specifically reducing *PVT1-MYC* but not canonical *MYC* expression (**Figures 4F, S4E and Table S4**). Together, our data suggest that *PVT1-MYC* fusion exhibits enhanced *MYC* oncogenic activity, contributing to its more potent role in cancer cell survival and gene regulation.

## Discussion

### EcDNAs are platforms for gene fusion

EcDNAs are pernicious driver of tumor evolution, acting as a platform for massive oncogene expression and rapid genome alterations. Uneven inheritance of ecDNA leads to intratumoral genetic heterogeneity and rapid tumor evolution. Because ecDNAs are themselves subject to sequence evolution with selection for contents that enhance tumor fitness^38,47,48,79^, ecDNA represents a powerful form of genome instability that generates fusion genes with transcripts that drive tumor growth. By integrating WGS and RNA-seq data in TCGA and CCLE databases, here we demonstrate that genomic rearrangements with high SV burdens on ecDNA drive amplified expression of gene fusions in cancers. Notably, oncogenes amplified on ecDNA emerge as hotspots for SVs, exhibiting heightened expression of oncogene fusion transcripts. Among these, *PVT1* is the most recurrent fusion oncogene across ecDNA(+) cancers, showing significant enrichment on ecDNA compared to other types of copy number variation. Our analysis reveals that *PVT1* fusions are dominantly enriched at the 5′ end to include its promoter and exon 1. We find fusion of *PVT1* exon 1 to 5’ of oncogenes or reporter genes confers increased mRNA stability. Using the most prevalent *PVT1* fusion in cancer–*PVT1-MYC* as a model and leveraging isoform-specific single-cell RNA-seq, we demonstrate that *PVT1* fusion enhances oncogenic function of *MYC* by inducing its downstream target genes. This study uncovers a previously unrecognized link between ecDNA and gene fusion and provides mechanistic insights into the prevalence and oncogenic role of *PVT1* fusion in cancer (**Figure 4G)**.

The strong association between *PVT1* fusion and ecDNA likely arises from both biological and structural factors. *PVT1* was first identified as a breakpoint hotspot in Burkitt’s lymphoma^80–82^, highlighting its fragility to genomic rearrangements. Given the heightened DNA damage response and repair activity on ecDNA and presence of multiple ecDNA copies in ecDNA hubs^4,12,83–86^, the genomic fragility of the *PVT1* locus may lead to increased fragmentation and reassembly on ecDNA, driving the amplification of *PVT1* fusions in cancer. In line with this, in COLO320DM ecDNA, *PVT1-MYC* fusions exist in three copies with a specific orientation^4,87^, suggesting that their formation involves multiple fragmentation and reassembly events rather than a single and random genomic rearrangement. EcDNA may serve as a platform for this stepwise, iterative process, facilitated by its enriched environment of elevated DNA damage response and repair activity. Beyond its structural instability, selection pressure on ecDNA may further reinforce the prevalence of oncogenic *PVT1* fusion in cancer. Similarly, EGFRvIII mutants, which are constitutively active due to the loss of the ATP-binding domain, become dominant when amplified on ecDNA in glioblastoma^8,88^. Once *PVT1* fusion is acquired by ecDNA, it may be selectively retained to preserve functionally advantageous fusion genes, further strengthening the link between ecDNA and oncogenic gene fusions. Future studies understanding the mechanisms driving *PVT1* fusion formation and selection on ecDNA could provide critical insights into cancer evolution and inform therapeutic strategies targeting ecDNA-driven oncogenic fusions.

### Oncogene activation by RNA stabilization

These results highlight an unexpected consequence of gene fusion on post-transcriptional RNA regulation. Gene fusions in cancer have previously been considered in the context of altered transcriptional regulation (e.g. *TMPRSS2-ERG*)^89–91^ or altered protein products (BCR-ABL)^92,93^. Recognition of their roles in cancer have led to improved cancer diagnostic strategies and targeted therapies. Our studies of *PVT1* 5’ fusions indicate that *PVT1* exon 1 serves as a universal 5’ UTR that helps to stabilize otherwise unstable oncogenic mRNAs, such as that encoding MYC. *PVT1-MYC* likely emerged as a frequent product both because of the proximity of the two fusion partner genes in the endogenous locus and resulting ecDNAs, as well as short half-life of *MYC* mRNA that makes the *PVT1* boost critical to extend the oncogene mRNA function. Endogenous *PVT1* is a chromatin-associated lncRNA^94^, and its fusion and export into the cytoplasm may liberate *PVT1* exon 1 from nuclear RNA decay pathways that would normally ensure its destruction. It will be interesting to study when and how SRSF1 binds to *PVT1* exon 1 and functions to increase RNA stability in the future studies.

Furthermore, cancer cells may take advantage of *PVT1* 5’ end fusion to boost *MYC* oncogene expression by replacing the *MYC* promoter with *PVT1* promoter^4^, which has been shown to compete with *MYC* promoter for enhancer binding^52^. In line with this, our data showed that in combination with *PVT1* promoter, *PVT1* exon 1 fusion synergistically increases RNA expression and enhances *MYC* rescue efficiency (**Figures 2G, S2C, and 4B**). *PVT1* promoter also facilitates episomes to join ecDNA hubs^4^. In normal cells, *MYC* expression can negatively regulate its own transcription^52,95^. However, cancer cells may evolve strategies to circumvent this autoregulatory feedback, and the *PVT1-MYC* fusion on ecDNA could play a role in disrupting this regulatory mechanism, contributing to tumor progression. Detailed mechanisms of fusion RNA splicing, transport, translation should be addressed in future studies.

Our study highlights the clinical potential of fusion RNAs as biomarkers in ecDNA-positive cancers. Early cancer detection is crucial for improving patient survival, yet reliable biomarkers remain limited. ecDNA can be detected in the precancerous stage and develops during cancer progression, increasing its structural complexity and copy number^11^, underscoring an urgent need for effective early detection strategies targeting ecDNA. Fusion RNAs derived from gene fusions amplified on ecDNA hold significant promise for enhancing cancer prognosis with greater sensitivity and accuracy. Combining *PVT1* fusion RNA detection with copy number variation analysis improves ecDNA detection accuracy to 95%, compared to 82.1% with copy number amplification (CN>10) alone or 74.5% with *PVT1* fusion RNA alone (**Figure S1I**). Furthermore, we provide a comprehensive list of genes amplified on ecDNA generating fusion RNAs, along with their prevalence in ecDNA-positive cancers, offering a new avenue for developing novel diagnostic tools with significant potential for improving ecDNA-related cancer prognosis. Importantly, detection of fusion transcripts offers greater clinical feasibility, given that SV analysis requires high sequencing depth, which challenges its clinical practice due to the high cost of data generation and storage.

### Limitations of the study

In this study, we focused on the fusion transcripts resulting from genomic rearrangements. However, fusion transcripts can also arise through genomic rearrangement-independent mechanisms such as trans-splicing between separate precursor mRNAs or alternative splicing of readthrough transcripts^21,96^. Our *MYC* ChIRP-MS analysis identified SR family proteins as being enriched in COLO320DM, which are known to be splicing factors, and we demonstrated that SRSF1 binding to *PVT1* exon 1 plays a role in *PVT1* exon 1-mediated mRNA stabilization (**Figure 3**). Additional studies are needed to further elucidate the molecular mechanisms on how SRSF1 binding stabilizes mRNA and to explore the interplay between splicing and mRNA stability. Additionally, we used a naturally occurring cell line model harboring *PVT1-MYC* genomic rearrangement on ecDNA, which represents the most prevalent form of gene fusion associated with ecDNA. Future studies will be needed to understand the full landscape of gene fusions arising on ecDNA, their functional impact, and how they contribute to tumor growth and treatment resistance employing ecDNA engineering.

**Figure S1.**
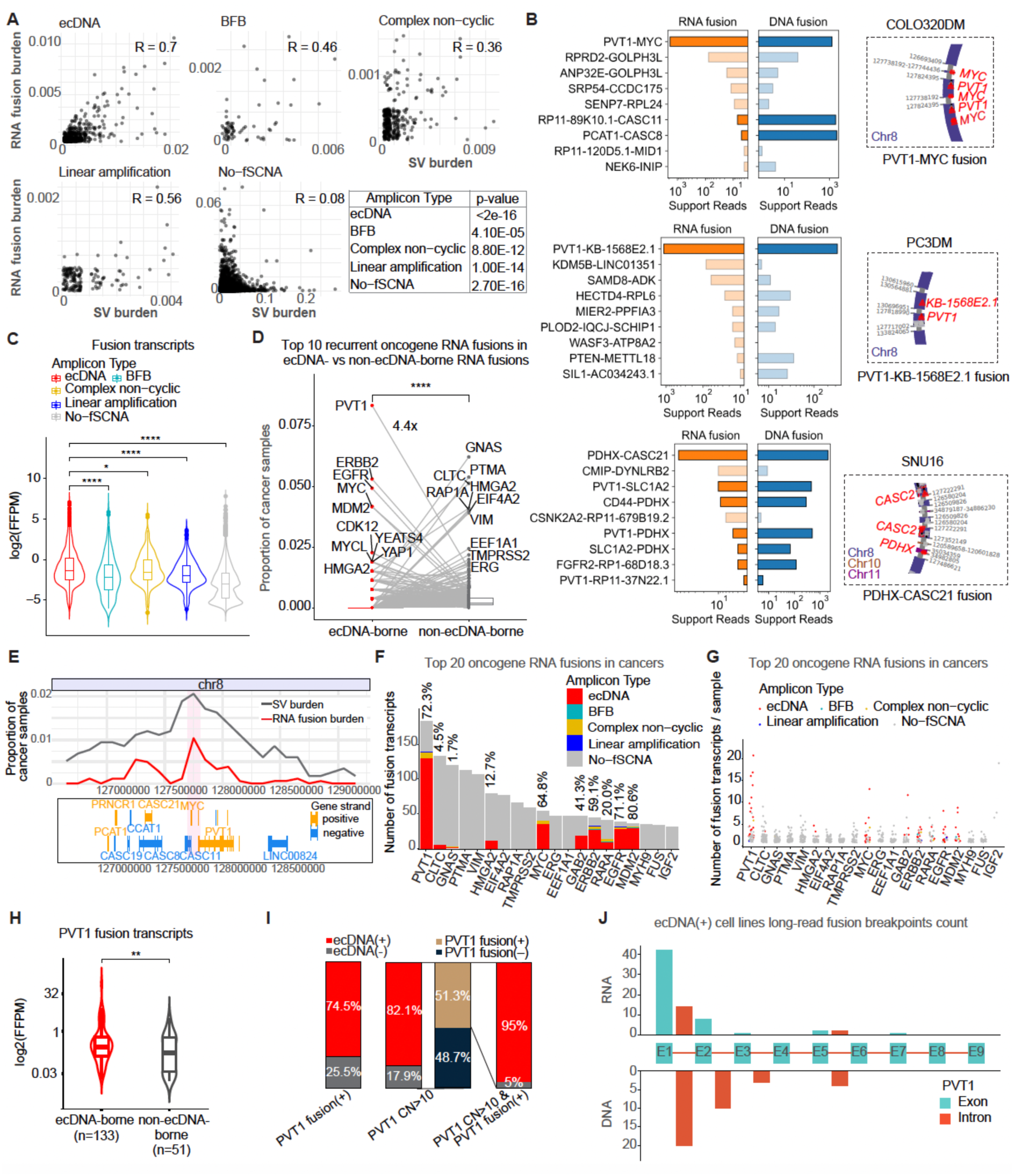
Landscape of gene fusions on ecDNAs, related to Figure 1. (A) Correlation between SV and RNA fusion burden by amplicon type (TCGA and CCLE). Each dot represents a 100 kb genomic window, categorized by amplicon type. Pearson’s correlation coefficient (R) is denoted within each plot, and p-values were determined by two-tailed unpaired t-test. (B) Number of RNA fusion reads (left) and DNA fusion reads (middle), detected by long-read sequencing for genes expressing the top 9 most abundant fusion transcripts in ecDNA(+) cell line models. Fusion genes amplified on ecDNA are presented as dark colored bars while the bars of non-ecDNA genes are light-colored. The right panel shows the genomic structures of ecDNA supporting the most abundant RNA fusions, annotated with genomic coordinates based on CoRAL, a long-read sequencing-based ecDNA reconstruction method. (C) Expression level (log2FFPM) of fusion transcripts by amplicon type (TCGA and CCLE) (*p < 0.05, ****p < 0.0001; two-tailed unpaired t-test). (D) Proportion of cancer samples with oncogene RNA fusions, categorized by ecDNA-borne (red) vs non-ecDNA-borne RNA fusions (gray). The top 10 recurrent oncogenes with RNA fusions in each group are labeled, with lines connecting shared oncogenes between the two groups (****p < 0.0001; two-tailed unpaired t-test). (E) Distribution of SV and RNA fusion burden peaks near the *MYC/PVT1* region in ecDNA(+) cancers (TCGA and CCLE). SV and RNA fusion burden are defined as in Figure 1D. (F) Number of fusion transcripts for the top 20 oncogene RNA fusions across cancer samples (TCGA and CCLE). Amplicon types are color-coded in bars, with the percentage of ecDNA-borne RNA fusions indicated at the top of each bar. (G) Number of fusion transcripts per cancer sample for the top 20 oncogene RNA fusions shown in Figure S1F (TCGA and CCLE). Each dot represents a cancer sample harboring the corresponding oncogene RNA fusion, with amplicon type color-coded. (H) Expression level (log2FFPM) of *PVT1* fusion transcripts derived from ecDNA-borne and non-ecDNA-borne RNA fusions (TCGA and CCLE). The number of *PVT1* fusion transcripts used in the comparison is indicated in parentheses (**p < 0.01; two-tailed unpaired t-test). (I) Proportion of cancer samples with ecDNA (red bar) among *PVT1*-related cancers in three classifications (TCGA and CCLE): cancers with *PVT1* fusion transcripts (PVT1 fusion (+)), cancers with *PVT1* copy number variation (CN > 10), and cancers with both *PVT1* CN > 10 and *PVT1* RNA fusion. (J) Distribution of RNA fusion breakpoints (top) and DNA breakpoints (bottom) of *PVT1* fusions across exonic and intronic regions of *PVT1*, measured by long-read sequencing in ecDNA(+) cell line models and their isogenic pairs where *PVT1* fusion transcripts are detected (COLO320DM, COLO320HSR, PC3DM, SNU16, GBM39KT, and GBM39HSR).

**Figure S2.**
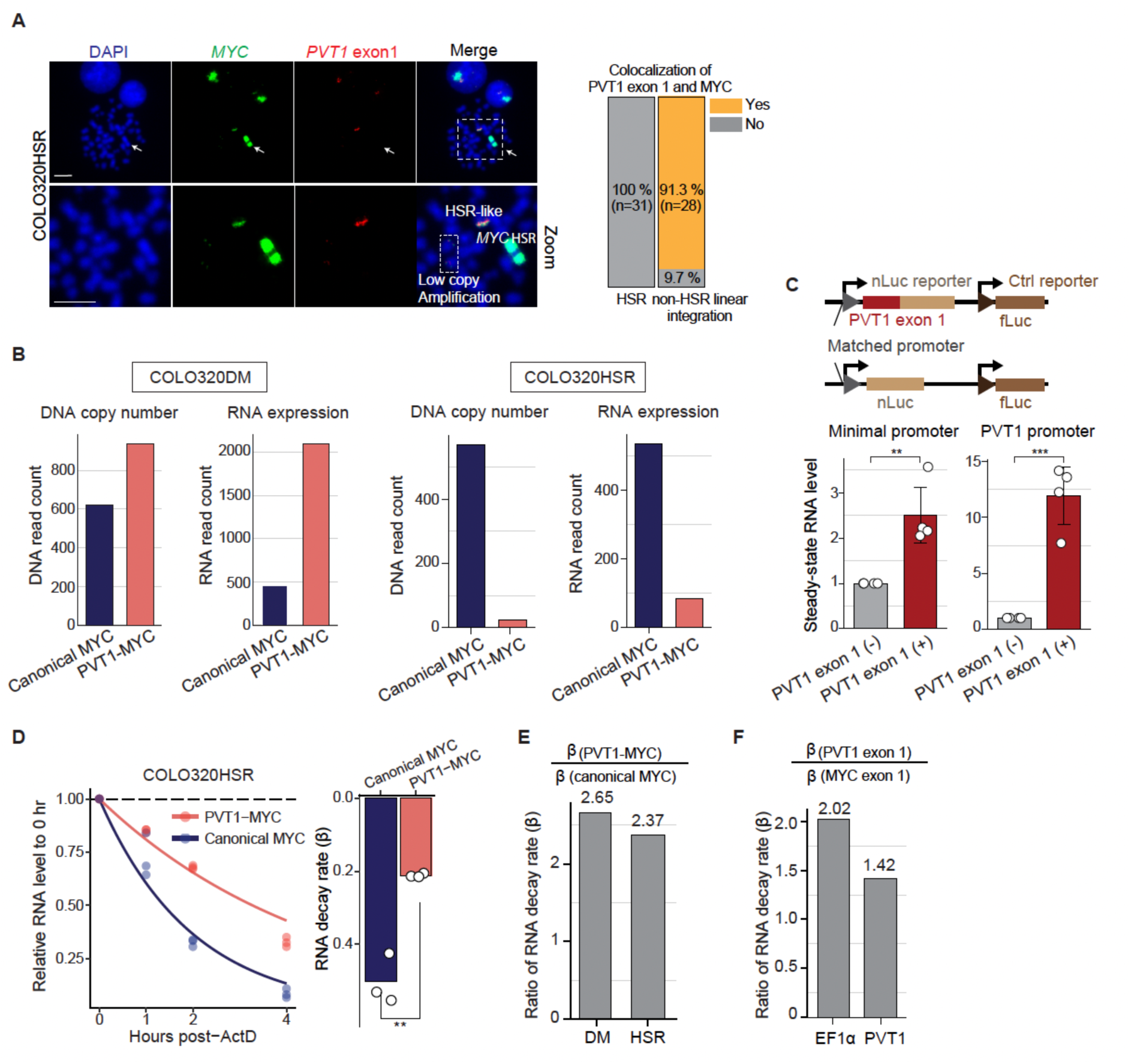
*PVT1* exon 1 fusion enhances mRNA stability, related to Figure 2. (A) Left: Magnified DNA FISH images of the COLO320HSR cell line in Figure 2B. Right: Quantification of colocalized signals between *PVT1* exon 1 and *MYC* on HSR and non-HSR linear integration from DNA FISH images of COLO320HSR cell line (n=31). (B) Raw DNA and RNA read counts of *PVT1-MYC* and canonical *MYC*, measured by long-read sequencing in Figure 2C–D. (C) Top: Schematic of the reporter assay system either with or without *PVT1* exon 1 fusion. Bottom: Steady-state RNA levels of nLuc reporter transcripts driven by either the minimal or *PVT1* promoter, normalized to fLuc internal control RNA (n=4). RNA levels were measured by RT-qPCR 48 hr post-transfection with equal amounts of plasmid DNA in COLO320DM cells (**p < 0.002, ***p < 0.0002; two-tailed t-test). (D) RNA stability and decay rate analysis of endogenous *PVT1-MYC* and canonical *MYC* transcripts in COLO320HSR cells using RT-qPCR. Left: Each dot represents relative RNA abundance normalized to 0 hr and *GAPDH* internal control RNA, with fitted lines based on decay rates from mathematical modeling. Right: RNA decay rate of canonical *MYC* and *PVT1-MYC*. Each dot represents a decay rate for each replicate (**p < 0.002; two-tailed t-test). (E) Fold change of RNA decay rates between *PVT1-MYC* and canonical *MYC* in COLO320DM and COLO320HSR cell lines in Figures 2F and S2D. (F) Fold change of RNA decay rates between *PVT1* exon 1- or *MYC* exon 1-fused reporters in Figure 2H.

**Figure S3.**
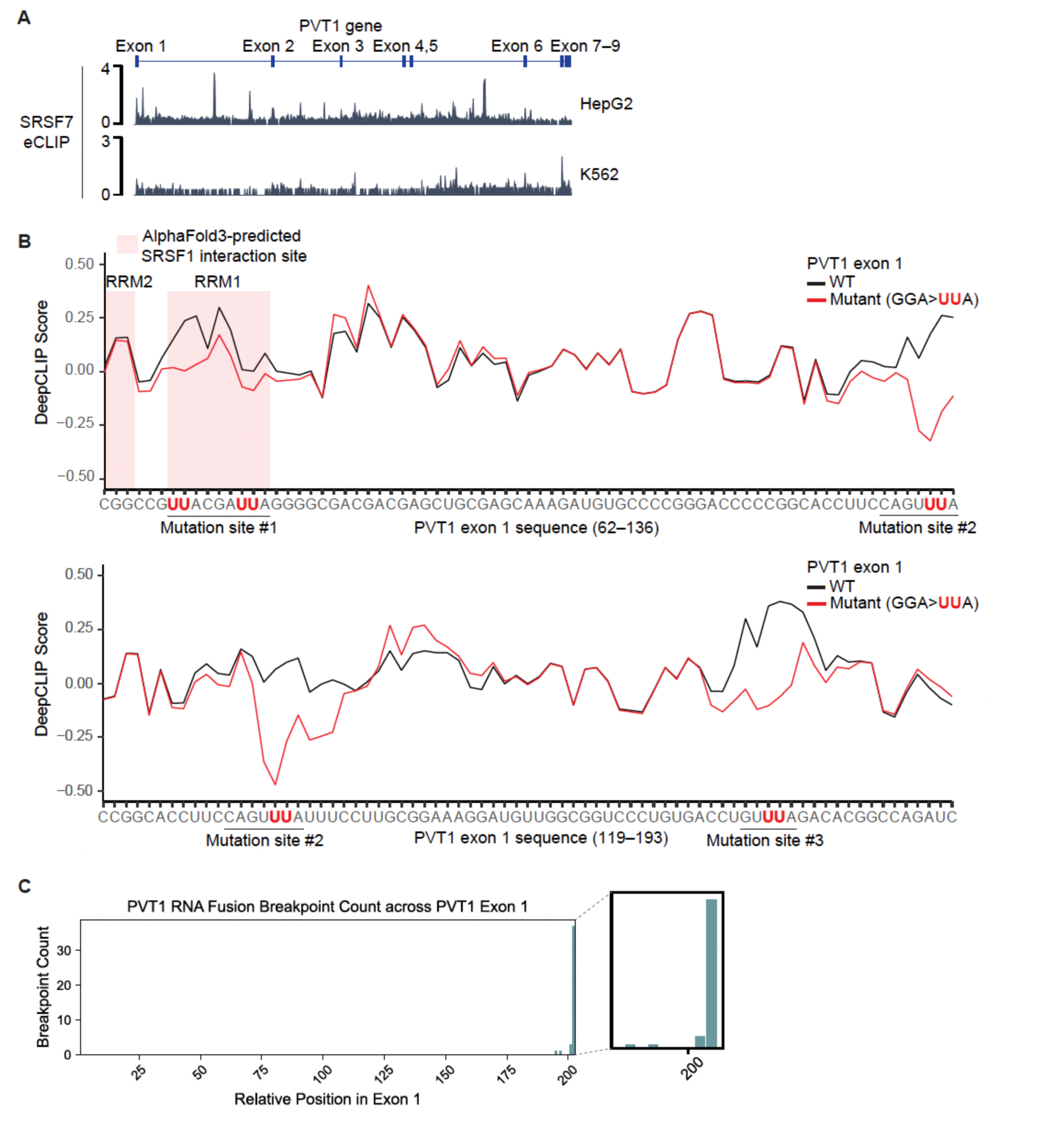
SRSF1 binding sites in *PVT1* exon 1, related to Figure 3. (A) Read density tracks of SRSF7 eCLIP across exonic and intronic regions of *PVT1* in HepG2 and K562 cell lines. (B) DeepCLIP score distribution along *PVT1* exon 1 mRNA sequence comparing the wild-type and mutant sequences based on ENCODE-K562 SRSF1 eCLIP data. A 75-nucleotide segment from each wild-type (WT) and mutant sequence containing the mutation sites was used as input for DeepCLIP score calculation. GGA > UUA substitutions in the mutation sites (underlined) are denoted in red. (C) Distribution of RNA fusion breakpoints across *PVT1* exon 1, measured by long-read RNA sequencing in ecDNA(+) cell line models and their isogenic pairs where *PVT1* fusion transcripts are detected (COLO320DM, COLO320HSR, PC3DM, SNU16, GBM39KT, and GBM39HSR).

**Figure S4.**
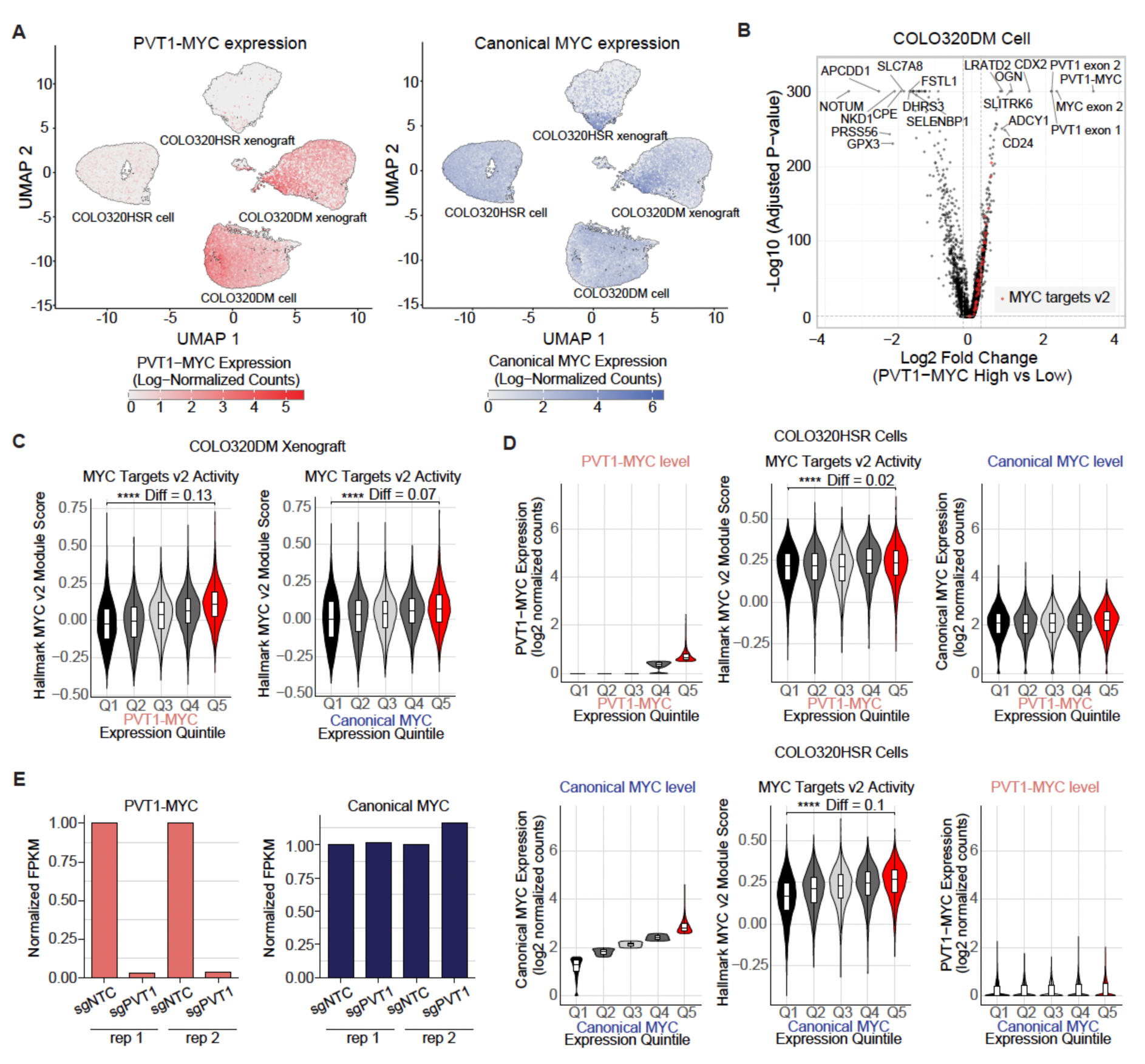
*PVT1-MYC* activates MYC targets, related to Figure 4. (A) UMAP visualization of single cells analyzed by Flex scRNA-seq in COLO320DM and COLO320HSR cells and xenograft mice *in vivo* models. Single-cell expression levels of *PVT1-MYC* (left) and canonical *MYC* (right) are shown as red and blue, respectively. Each dot represents a single cell. (B) Differentially expressed genes in *PVT1-MYC* high vs low COLO320DM cells, with the top 10 upregulated and downregulated genes labeled. MYC targets v2 genes are highlighted as red dots. (C) Violin plots displaying dose-dependent MYC targets v2 activity across expression quintiles of *PVT1-MYC* (left) and canonical *MYC* (right) in COLO320DM xenograft. Difference between Q5 mean and Q1 mean is presented as Diff (****p < 0.0001; Wilcoxon rank-sum-test). (D) Violin plots displaying dose-dependent MYC targets v2 activity and expression level of *MYC* isoforms across expression quintiles of *PVT1-MYC* (top) and canonical *MYC* (bottom) in COLO320HSR cells. Difference between mean of the highest quintile (Q5) and lowest quintile (Q1) is presented as Diff (****p < 0.0001; Wilcoxon rank-sum-test). (E) Normalized RNA expression levels of *PVT1-MYC* and canonical *MYC* in CRISPRi sgPVT1 RNA-seq in COLO320DM cells, relative to sgNTC (non-targeting control).

## Resource availability

### Lead contact

Further information and requests for resources and reagents should be directed to and will be fulfilled by the lead contact, Paul S. Mischel and Howard Y. Chang.

## Materials availability

This study did not generate new unique reagents.

## Data and code availability

High-throughput sequencing data from long-read DNA and RNA sequencing, Flex scRNA-seq, and CRISPRi PVT1 RNA-seq generated in this study have been deposited in the GEO and are publicly available as of the date of publication. This paper also analyzes existing, publicly available data. Accession numbers for all datasets are listed in the Key resources table. Custom codes used in this study will be made available on Github (https://github.com/ShuZhang0917/ecDNA-borne_fusion.git). Source imaging data generated for this study have been deposited in the Stanford Digital Repository (https://doi.org/10.25740/cp783mb3217).

## Acknowledgement

We thank members of the Chang and Mischel labs for discussion. This work was delivered as part of the eDyNAmiC team supported by the Cancer Grand Challenges partnership funded by Cancer Research UK (CGCATF-2021/100012 (P.S.M. and H.Y.C.), CGCATF-2021/100025 (V.B.), and the National Cancer Institute (OT2CA278688 (P.S.M. and H.Y.C.), OT2CA278635 (V.B.). L.A.G. is funded by the Arc Institute, NIH (UM1HG012660) and CRUK/NIH (OT2CA278665 and CGCATF-2021/100006). H.Y. is a Howard Hughes Medical Institute Fellow of the Life Sciences Research Foundation. Y.W. is supported by Schmidt Science Fellows program. K.L.H. was supported by a Stanford Graduate Fellowship and a NCI Predoctoral to Postdoctoral Fellow Transition Award (NIH F99CA274692). M.G.J. is supported by a NCI Pathway to Independence Award (NIH K99CA286968). A.B.-S. is supported by the Stanford Medical Scholars Research Program and in part by an Alpha Omega Alpha Carolyn L. Kuckein Student Research Fellowship. X.Y. is a Damon Runyon Fellow supported by the Damon Runyon Cancer Research Foundation (DRG-2474-22). J.A.B. is a Hanna Gray Fellow and H.Y.C. was an Investigator of the Howard Hughes Medical Institute. This work was supported by the Vincent Coates Foundation Mass Spectrometry Laboratory, Stanford University Mass Spectrometry (RRID:SCR_017801) utilizing the Bruker timsTOF Ultra & nanoElute 2 system (RRID: SCR_025639) purchased with generous support from Stanford C-ShaRP. This work was supported in part by NIH P30 CA124435 utilizing the Stanford Cancer Institute Proteomics/Mass Spectrometry Shared Resource.

## Author contributions

H.Y. and H.Y.C. conceived the project. H.Y., S.Z., P.S.M., and H.Y.C. designed the study. H.Y. conducted long-read RNA sequencing of all tested cell lines and DNA sequencing of COLO320DM and COLO320HSR cell lines. H.Y. conducted reporter construction, reporter assays for RNA regulation, ChIRP-MS, total MS and Flex scRNA-seq probe design. S.Z. analyzed AmpliconArchitect and SV data from WGS and RNA fusion data from RNA-seq in TCGA and CCLE databases and CRISPRi PVT1 RNA-seq. J.S. designed Flex scRNA-seq probes, generated and analyzed Flex scRNA-seq data. Y.W. generated long-read DNA sequencing of PC3DM cell line and analyzed long-read sequencing and public eCLIP data. V.P.K. performed and analyzed MYC rescue experiment. K.L.H analyzed mathematical modeling of RNA decay rate. I.T.-L.W. performed metaphase DNA FISH. S.S. performed AlphaFold3-based structural modeling of PVT1 exon 1 mRNA and SRSF1 RRM1–2 and analyzed DeepCLIP score. E.J.C. generated COLO320DM and COLO320HSR xenograft mice models. A.B.-S., K.K. and Q.S. generated CRISPRi PVT1 COLO320DM cell line, R.L. generated CRISPRi PVT1 RNA-seq and X.Y. generated COLO320DM dCas9-BFP-KRAB (CRISPRi) cell line. M.G.J. analyzed the long-read RNA-seq. J.L. provided AmpliconArchitect results from TCGA and CCLE databases. Y.Z. provided TCGA SV call results from GDC. S.K.P. cloned luciferase reporter constructs under H.Y.’s guidance. F.M.L. helped structural modeling analysis under S.S.’s guidance. H.Y., S.Z., K.L.H., J.A.B., V.B., D.W.F, L.G., P.S.M, and H.Y.C. guided data analysis and provided feedback on experimental design. H.Y., S.Z., P.S.M, and H.Y.C. wrote the manuscript with input from all the authors. H.Y.C. and P.S.M supervised the project.

## Methods

### Key resources table

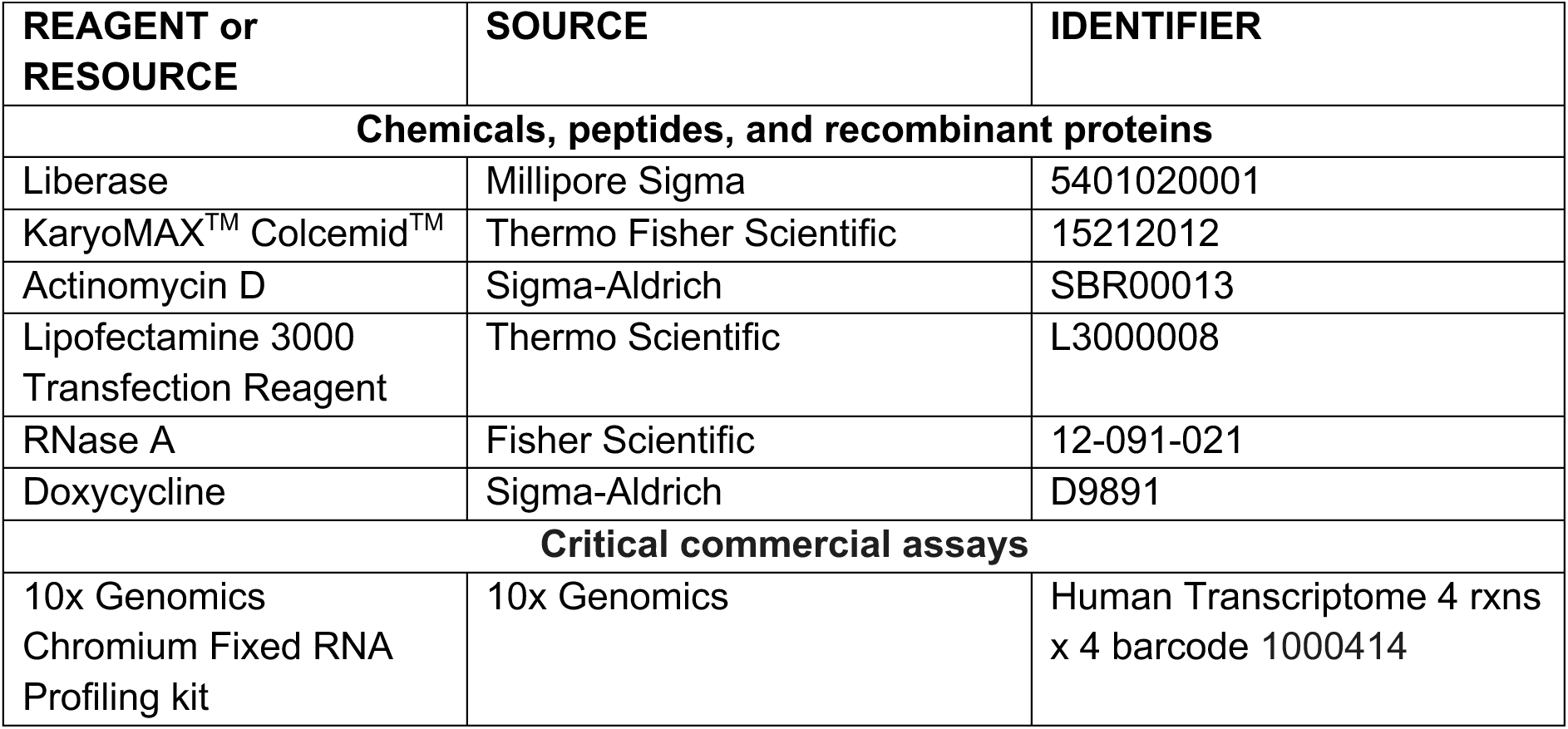
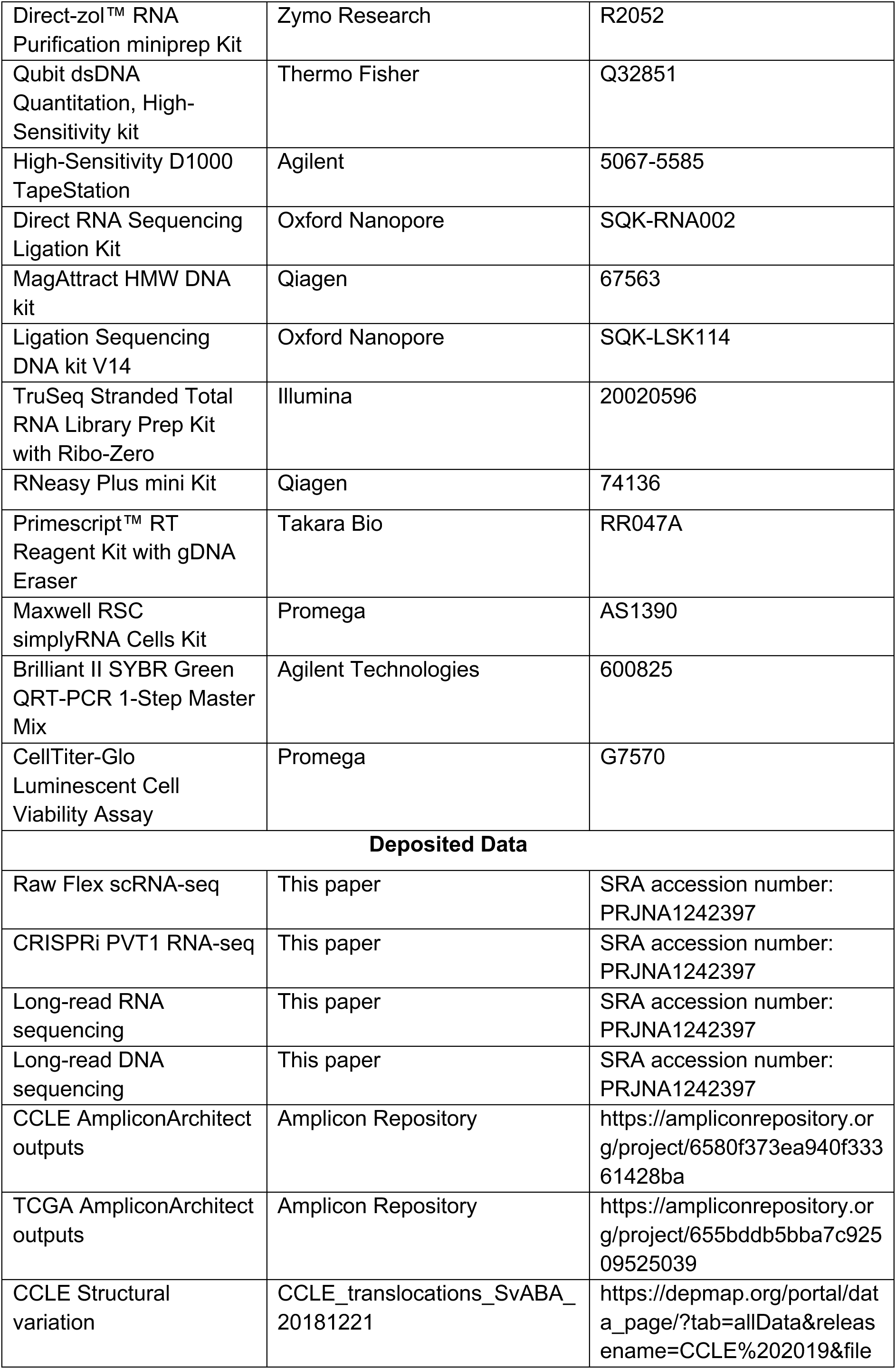
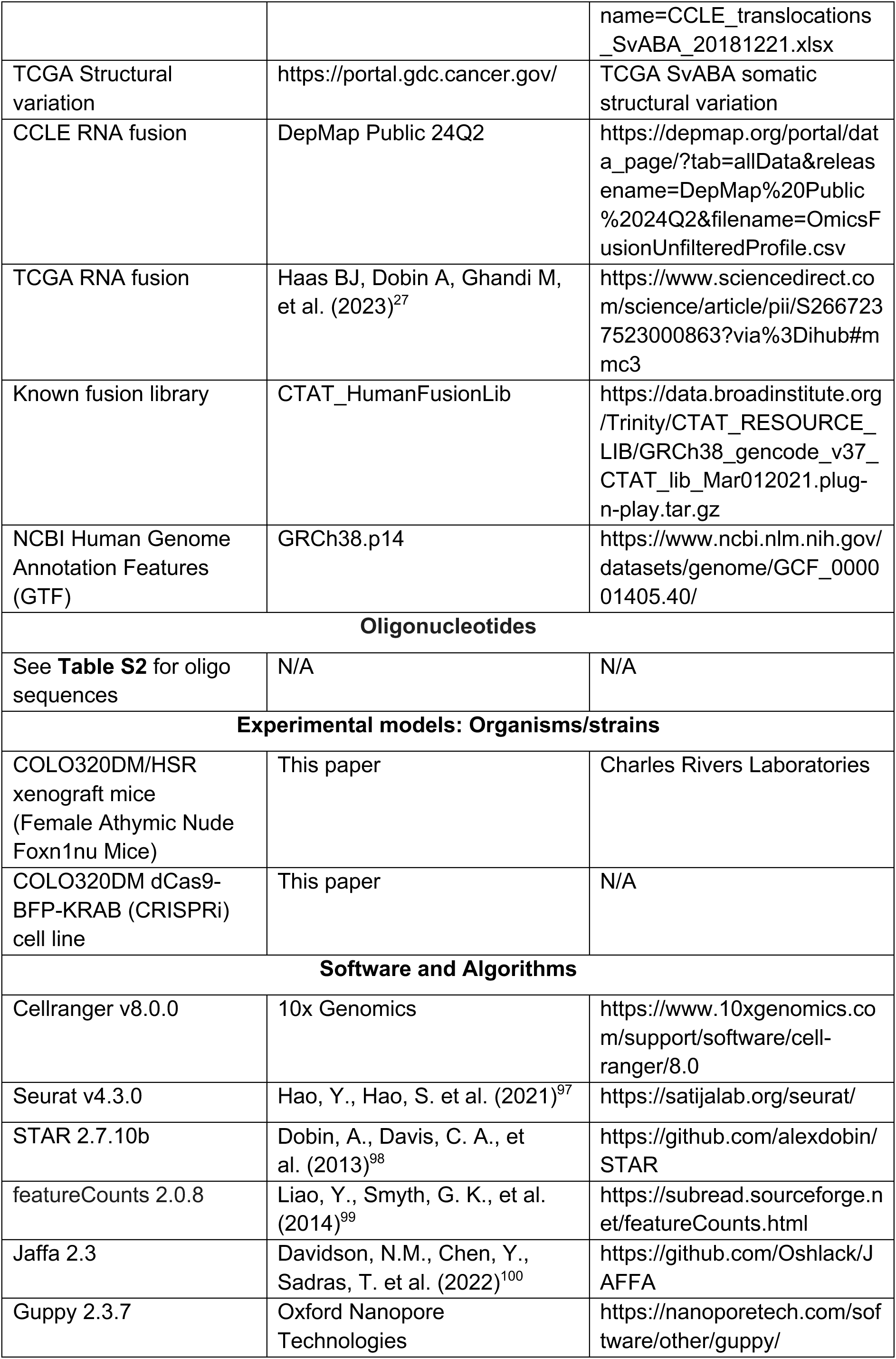
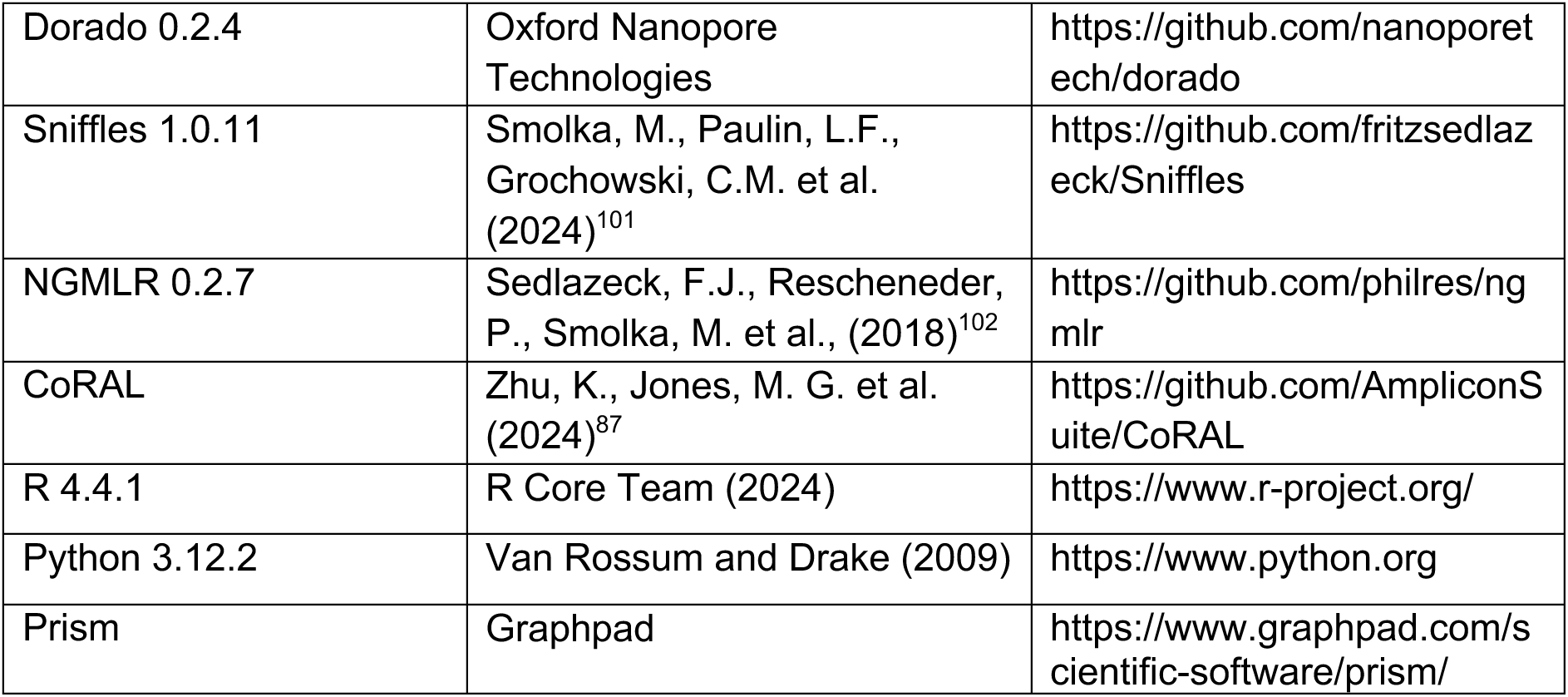

### Cell culture

The parental PC3 line was obtained from ATCC. PC3DM line was isolated by the Mischel Lab through single-cell expansions of the parental PC3 line^85^. All the other cell lines were purchased from ATCC. Human colorectal cancer cell line COLO320DM, COLO320HSR, and HCT116; prostate cancer cell line PC3DM; gastric cancer cell line SNU16; embryonic kidney cell line HEK293T; and hTERT-immortalized retinal pigment epithelial cell line RPE1 were cultured in 4.5 g l^−1^ glucose-formulated Dulbecco’s Modified Eagle’s Medium (DMEM; Thermo Fisher Scientific, 11995073) supplemented with 10% fetal bovine serum (FBS; Gibco) and 1% penicillin–streptomycin (pen-strep; Thermo Fisher Scientific, 15140163). GBM39 cell lines were cultured in Dulbecco’s Modified Eagle’s Medium/F12 (Gibco, 11320-033) supplemented with 1× B27 (Gibco, 17504-01), 20 ng ml^−1^ epidermal growth factor (Sigma, E9644), 20 ng ml^−1^ fibroblast growth factor (Peprotech, AF-100-18B), 1–5 µg ml^−1^ heparin (Sigma, H3149), 1× GlutaMAX (Gibco, 35050-061) and 1% pen-strep. All the cells were maintained at 37 °C with 5% CO2 in a humidified incubator. Cell lines routinely tested negative for mycoplasma contamination.

### RNA Fusion analysis using TCGA and CCLE databases

We applied a comprehensive filtering strategy for unfiltered RNA fusions identified by STAR-Fusion within the CCLE and TCGA RNA-seq datasets. Specifically, CCLE RNA fusions from 1,482 cell lines were sourced from the DepMap 24Q2 Public dataset (OmicsFusionUnfilteredProfile.csv), while TCGA RNA fusions, encompassing 9,426 tumor samples and 707 normal samples, were extracted from Table S2 of Brian J. Haas et al. Cell Reports Methods 2023^27^.

To minimize potential false positives, we implemented several exclusion criteria based on previous studies^27,29,103^: (1) Red Herrings: Fusion transcripts classified as Red Herrings by FusionInspector (version 2.8.0) were excluded based on known fusions derived from normal samples, as specified in GRCh38_gencode_v37_CTAT_lib_Mar012021.plug-n-play.tar.gz. (2) Gene Origin: Fusion partners that originated from the same gene or included paralogous genes, as annotated in the GRCh38.p14 GTF file and accessed through Ensembl Biomart, were discarded. (3) Normal Fusions: We filtered out common fusion transcripts identified in cancer samples that were recurrently observed in TCGA normal samples. (4) Gene Types: Fusion partners involving mitochondrial genes, HLA genes, or immunoglobulin genes were also removed from our analysis. (5) Expression Thresholds: Non-recurrent fusions with low expression levels, defined as a Fusion Fragments Per Million (FFPM) < 0.1, were excluded. Additionally, recurrent fusions with a maximum FFPM less than 0.05 were filtered out. Furthermore, we removed highly recurrent fusions that were exclusive to CCLE, characterized by the same breakpoints in at least 10 samples, as long as they were not present in TCGA tumor samples.

### Amplicon type analysis

AmpliconArchitect (AA) was utilized to identify focally amplified regions, which were classified into four distinct types: extrachromosomal DNA (ecDNA), breakage-fusion-bridge (BFB) cycles, complex non-cyclic amplifications, and linear amplifications. Genomic regions without any detected focal amplifications were defined as no focal somatic CN amplification (No-fSCNA^5^). The AmpliconArchitect results were obtained from the repository at https://ampliconrepository.org/^37^, where we downloaded data for 329 CCLE cell lines and 2,471 TCGA samples.

For the structural variants (SVs), we retrieved CCLE SV data for 329 cell lines from DepMap (CCLE_translocations_SvABA_20181221)^104^. TCGA somatic SVs from 844 tumor samples that overlapped with AmpliconArchitect samples were acquired from the GDC (Genomic Data Commons) data portal at https://portal.gdc.cancer.gov/. Both CCLE and TCGA SVs were identified using the SvABA tool.

To ensure the integration of amplicon types, SVs, and RNA fusions that were originally aligned to different versions of the human genome, we converted the TCGA AmpliconArchitect results and CCLE SVs from GRCh37 to GRCh38 using the Liftover tool. Then, we aligned TCGA Tissue Source Site and the participant number to match RNA-seq data to the WGS data from the same patient. Subsequently, we annotated the genes associated with the focal amplicons as well as the nearest genes within a 100 kb range, both 5’ and 3’ of the SV breakpoints, utilizing the GRCh38.p14 GTF file. To identify SVs supporting gene fusions (SVGF) producing fusion transcripts, SV breakpoints were assigned if they were less than 500 kb from RNA fusion breakpoints and within 100 kb of a gene body^20^. Finally, if the RNA fusion breakpoints and SV breakpoints were located within amplicons that were predicted as specific amplicon types, we categorized those breakpoints accordingly. This classification allowed us to associate each breakpoint with its respective amplicon type, facilitating further analysis to compare the features of SVs and RNA fusions from different amplicon types.

### Analysis of fusion rate and burden for SVs and RNA Fusions

Using the matched samples with AA, SV and RNA fusion data, the rate for SVGF in each amplicon interval was calculated as dividing the number of genes with SV breakpoints within or near (<100 kb) the gene body^44–46^, by the total number of genes within each amplicon interval. The RNA fusion rate was determined as the number of gene with RNA fusion breakpoints within the gene body divided by the total number of gene within each amplicon interval.

SV burden^19^ and RNA fusion burden were calculated as the proportion of cancer samples harboring SV or RNA fusion breakpoints within 100 kb genomic windows across entire genome. The number of samples with SV or RNA fusion breakpoints were divided by the total number of samples linked to each amplicon type.

### Long-read RNA sequencing

Total RNA was extracted by Direct-zol™ RNA Purification miniprep Kit (Zymo Research, R2052). RNA integrity (RIN) was confirmed by Bioanalyzer and RNA samples with maximum RNA integrity (RIN = 10) were used for Nanopore library preparation. We constructed Nanopore libraries using the Oxford Nanopore Direct RNA Sequencing Ligation Kit (SQK-RNA002) according to manufacturer’s instructions. We sequenced libraries on an Oxford Nanopore PromethION using a R.9.4 Flow Cell (FLO-PRO002) according to manufacturer’s instructions.

To identify fusion transcripts, we utilized Jaffa^105^ (version 2.3) on our Oxford Nanopore Direct RNA-sequencing (DRS) data. Specifically, basecalls were derived from raw FAST5 signal using Guppy (using the 2020-09-21_rna_r9.4.1_promethion_256_855130ab RNA basecalling model). Then, Jaffa was invoked using the default .groovy pipeline with the following flags: -n 20. The resulting Jaffa file was used to determine the support and confidence of each identified fusion transcript.

### Long-read DNA sequencing library preparation

High-molecular weight (HMW) genomic DNA of COLO320DM, COLO320HSR, PC3DM and SNU16 cells was extracted from approximately 2 million using MagAttract HMW DNA kit (Qiagen, 67563). After extracting HMW gDNA, we constructed Nanopore libraries using the Oxford Nanopore Ligation Sequencing DNA kit V14 (SQK-LSK114) according to manufacturer’s instructions. We sequenced libraries on an Oxford Nanopore PromethION using a 10.4.1. Flow Cell (FLO-PRO114M) according to manufacturer’s instructions. Basecalls from raw POD5 files were computed using Dorado (v.0.7.2). DNA reads were aligned to hg38 using minimap2 (v2.26).

To detect SVs that support RNA fusion breakpoints, we considered SV breakpoints within or near (<100 kb) the gene body^44–46^. DNA long read that support the RNA gene fusions in **Figure S1B** were identified based on the SA tag of minimap2 aligned reads. *PVT1* DNA breakpoint positions in **Figure S1J** were identified using Sniffles2 (v2.0.7). ecDNA structure was reconstructed by referring to Complete Reconstruction of Amplifications with Long reads (CoRAL) for reconstructing ecDNA architectures using long-read data^87^.

### Generation of xenograft mice model

Female athymic nude mice (Charles River Laboratories) were housed under standard conditions and used for subcutaneous tumor implantation. Prior to injection, mice were anesthetized using isoflurane in an induction chamber. A 100 µL aliquot of the prepared cell suspension was injected bilaterally into the subcutaneous space of the flanks using a 25-gauge needle and a 1 mL syringe. Mice were monitored post-procedure for signs of distress or complications.

At the experimental endpoint, mice were euthanized using a sealed CO₂ chamber, with cervical dislocation performed as a secondary confirmatory method. Tumors were sharply dissected from the subcutaneous space, immediately flash-frozen in liquid nitrogen, and stored at −80°C for future analysis.

### Fixed scRNA-seq (Flex-seq)

Xenografts were dissociated according to 10x Genomics Tissue Fixation and Dissociation Demonstrated Protocol #CG000553 (Rev B). Xenografts were stored in liquid nitrogen until fixation where they were thawed at room temperature and sectioned into ~25 mg tissue samples which were titurated with a wide-bore P1000 in fixation buffer incubated for 20 hours at 4 °C. Fixed tissue sections were quenched and supplemented with Enhancer to 10% and 50% w/v Glycerol to 10% and stored at −80 °C. Frozen fixed tissues were washed and resuspended in 0.2 mg/ml Liberase (Millipore Sigma 5401020001) prewarmed to 37 °C and incubated for 30 minutes, samples were titurated and dissociation was evaluated by Countess II FL Automated Cell Counter (Thermo Fisher Scientific) every 10 minutes.

Cell samples were fixed with Chromium Next GEM Single-Cell Fixed RNA Sample preparation kit (10x Genomics 1000414) was used according to the manufacturer’s protocol. Aliquots of 2 million cells were fixed according to the Fixation of Cells and Nuclei Demonstrated Protocol # CG000478 (Rev D). Fixation for 20 hours at 4 °C, was quenched and supplemented with Enhancer to 10% and 50% w/v Glycerol to 10% and stored at −80 °C. Cryopreserved fixed samples were thawed and ~500,000 cells were hybridized using Chromium Fixed RNA Kit, Human Transcriptome, 4rxn x 4BC (PN-1000475, 10x Genomics). LHS and RHS custom probe pools were added to hybridization reactions according to their barcode to a final concentration of 2 nM following 10x Genomics Demonstrated Protocol #CG000621 (Rev D). Hybridized cell and xenograft samples were pooled and washed following the Pooled Wash Workflow #CG000527. Two lanes of GEMs were generated on the Chromium X (10x Genomics) targeting 60,000 cells recovered.

Single-cell gene expression of COLO320DM and COLO320HSR cells and xenografts was assayed using 10x Chromium Fixed RNA-Profiling scRNA-seq. Indexed libraries were quantified by Qubit dsDNA Quantitation, High-Sensitivity kit (Thermo Fisher Q32851) for yield and High-Sensitivity D1000 TapeStation (Agilent 5067-5585) to ensure expected library size and absence of contaminating products. Pooled libraries were sequenced by paired-end: Read 1: 28, I1: 10, I2: 10, Read 2: 90 (Illumina Nova-Seq X).

### scRNA-seq analysis

BCL Convert Software (Illumina) was used to generate demultiplexed FASTQ sequencing libraries which were aligned with Cellranger multi (v.8.0.0, 10x Genomics) using a human probeset reference (v1.0.1-GRCh38-2020-A) modified to include custom probe sequence references (Sequences provided in **Table S2**). Seurat objects were constructed from the Cellranger generated filtered feature-barcode matrices and further filtered for cells with greater than 200 detected genes and less than 10% of mitochondrial reads^106^. Dimensionality reduction was performed on the top 2,000 most variable genes of log-normalized and scaled samples.

Expression quantiles for PVT1-MYC were generated by binning each sample by depth-normalized counts. MYC isoform driven differentially expressed genes were calculated on the output FindMarkers with a Wilcoxon rank-sum test comparing PVT1-MYC high (Q5) vs low (Q1) with a log2-foldchange threshold of 0.25. Gene-set enrichment on these differentially expressed genes were calculated by GSEA for Hallmark signatures. Cells were scored for the expression of Hallmark MYC v2 signature using the AddModuleScore function (Seurat) with 100 control genes. MYC v2 Hallmark module activity between PVT1-MYC quintiles was calculated by Wilcoxon rank-sum-test.

### DNA FISH staining and imaging analysis

COLO320DM and COLO320HSR cells were arrested in metaphase with 100ng ml^-1^ KaryoMAX^TM^ Colcemid^TM^ Solution in PBS (Gibco) for 4 hours. The cells were collected after trypsinization and washed once in PBS. The cells were resuspended in 0.075M KCl for a 20-min incubation at 37°C, followed by fixation with Carnoy’s fixative (3:1 methanol: acetic acid). The cell pellet was washed thrice with Carnoy’s fixative prior to being dropped onto a humidified glass slide. After the sample was fully air-dried, the slide was briefly incubated in 2X SSC buffer, followed by dehydration in ascending ethanol concentrations of 70%, 85% and 100% for 2 mins each. FISH probes obtained from Empire Genomics were freshly diluted in hybridization buffer at 1:6 ratio. The MYC region was probed with RP11-440N18 (Chr 8:128,596,756 – 128,777,986 - hg19). and PVT1 exon 1 region was targeted with the WI2 clone G248P89481C4 (Chr8:128,790,936-128,830,829 - hg19). The diluted probes were then added to the sample with a coverslip applied on top. The slide was subjected to heat denaturation at 75°C for 3 mins, followed by overnight incubation at 37°C for hybridization in a humidified slide moat. The next day, the coverslip was gently removed from the sample, and the slide was washed in 0.4X SSC and 2X SSC-0.1% Tween-20 for 2 mins each. DNA was stained with DAPI stain (50ng mL^-1^), followed by a brief wash in ddH2O and was left to air-dry. The sample was mounted with ProLong Diamond (Invitrogen). Image acquisition was performed on a Leica DMi8 widefield microscope with a 63x oil objective.

### RNA stability assay

For endogenous mRNA stability assay, 400,000 of COLO320DM and COLO320HSR cells were seeded in 12 well plate 1 day before the actinomycin D treatment. Next day, cells were treated with 4 μg/ml of actinomycin D and collected in a time course manner (0, 1, 2, and 4 hr). For reporter mRNA stability assay, 200,000 of COLO320DM cells were seeded in 12 well plate 1 day before plasmid transfection using the Lipofectamine 3000 Transfection Reagent (Thermo Scientific, L3000008) according to manufacturer’s instructions. Cells were treated with 4 μg/ml of actinomycin D 2 days post-transfection and collected in a time course manner (0, 2, 4 hr). For mutant reporter mRNA stability assays, 100,000 of HEK293T cells were seeded in 12 well plate 1 day before plasmid transfection and RNA stability assay was done as described above.

### Modeling RNA stability

To model the life cycle of RNA transcripts (Furlan et al. Briefings in Bioinformatics 2021^107^), we used an ordinary differential equation to represent the synthesis and decay of RNA,

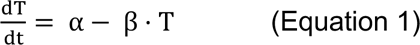

where T, the total number of RNA transcripts, changes over time t depending on the rates of synthesis (α) and decay (β). At steady state, the change in RNA transcripts over time is zero:

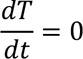

Therefore,

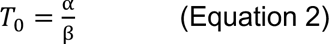

To calculate the RNA decay rate β, we analyzed quantitative reverse transcription PCR data for COLO320DM and COLO320HSR cells after treatment with actinomycin D at various timepoints. The total RNA transcripts at time t drop according to the following equation:

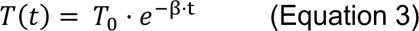

Where total RNA T(t) at time t is a function of total RNA at the start of transcription block, T0, the RNA decay rate β, and time t. This gives the RNA decay rate

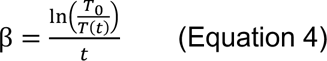

β was calculated for each transcript (canonical MYC and PVT1-MYC) for each experimental replicate in each cell line at each time point, and the mean β value of each replicate was calculated using all time points. Finally, to compare the contribution of RNA decay to total RNA transcript levels, we compared the RNA decay rates β of canonical MYC and PVT1-MYC with the total transcript levels per DNA copy at steady state using Equation 2.

### Reporter Plasmid Construction

To generate reporter constructs for testing PVT1 exon 1 fusion effect on RNA stability under the matched promoter–either PVT1 or control promoters (**Figures 2I and S2C**), sequence of PVT1 promoter (chr8:127793691–127794532, hg38), PVT1 exon 1 (chr8:127794533–127794734, hg38) and minimal promoter was amplified from the plasmid constructs (PVT1p-nLuc and minp-nLuc) used in Hung et al. Nature 2021^4^ and subcloned to reporter plasmids driving NanoLuc luciferase (nLuc) and a constitutive thymidine kinase (TK) promoter driving Firefly luciferase (fLuc) as an internal control by Gibson assembly.

To generate MYC-Flag-mNG11 reporter for testing PVT1 exon 1 fusion function under the matched promoter–either PVT1 or control promoters (**Figures 2I–J, S2D, 3F–G, 4A–B**), endogenous PVT1-MYC and canonical MYC sequence was amplified from cDNA that was reverse transcribed from total RNA of COLO320DM using Primescript™ RT Reagent Kit with gDNA Eraser (Takara Bio, RR047A) according to manufacturer’s instruction, and subcloned to a vector with Flag-mNG11 tag and a constitutive thymidine kinase (TK) promoter driving mCherry as an internal control.

### RT-qPCR

RNA was extracted using RNeasy Plus mini Kit (Qiagen, 74136) or Maxwell RSC simplyRNA Cells Kit (Promega, AS1390) with DNase treatment. 10 ng of RNA was used for RT-qPCR with 1× Brilliant II qRT-PCR mastermix with 200 nM forward and reverse primer and 0.5 μl RT/RNase block (Agilent, 600825) for 10 μl of total reaction. Each Ct value was measured using Lightcycler 480 (Roche) and each mean dCt was averaged from a duplicate RT–qPCR reaction with biological replicates. This list of RT-qPCR primers is shown in **Table S2**.

### ChIRP-MS

Cell harvesting, lysis, and ChIRP were performed largely as previously described^108^. Approximately 100 million cells were fixed with 3% formaldehyde for 30 min, followed by final 125 mM Tris-HCL pH 8.0 (Invitrogen, 15568-025) quenching for 5 min. Lysate was generated by resuspending cell pellets in 1 mL lysis buffer (50 mM Tris-pH 7.0, 10 mM EDTA, 1% SDS) per 100 mg of cell pellet weight. Sonication was done using a focused-ultrasonicator (Covaris E220) until the RNA length was ~500 nt as determined by agarose gel analysis and stored at −80°C. Lysates were thawed on ice and precleared by using 30 µL washed MyOne C1 beads (Thermo Fisher Scientific, 65002) per mL of lysate at 37°C for 30 min on rotation. Preclearing beads were removed twice from lysate using magnetic stand. For RNase control ChIRP, precleared lysates were treated with 30 µg of RNase A (Fisher Scientific, 12-091-021) per mL of lysate and incubated at 37°C for 45 min. Next, every 1 mL of experimental and RNase control ChIRP lysates were incubated with 100 pmole of biotinylated probes targeting *PVT1* exon 1 and *MYC* exon 1–3 with 2 mL of hybridization buffer (750 mM NaCl, 1% SDS, 50 mM Tris-HCl pH 7.0, 1 mM EDTA, 15% formamide) and incubated at 37°C for 16 hr on rotation. ChIRP probe pools were composed of an equimolar mix of antisense oligos (see **Table S2** for sequence). Next day, 100 µL of MyOne C1 beads for every 100 pmoles of probes (; per mL of lysate) were washed three times before use and incubated with lysates at 37°C for 45 min on rotation. RNA-protein interactome by ChIRP was collected on the beads with a magnetic stand and beads were washed 5 times for 2 min with constant mixing in 1 mL of ChIRP wash buffer (2x NaCl-Sodium Citrate (SSC, ThermoFisher Scientific), 0.5% SDS) at 37°C. At the last wash, 1% of beads were saved for RNA extraction for RNA pull-down QC. The enriched proteins were eluted by 600 µL of ChIRP biotin elution buffer (12.5 mM biotin, 7.5 mM HEPES, pH 7.9, 75 mM NaCl, 1.5 mM EDTA, 0.15% SDS, 0.075% sarkosyl, and 0.02% Na-Deoxycholate), on rotation at 25°C for 20 min and at 65°C for 15 min shaking. Eluent was transferred and pooled by total twice elution (~1200 μL), and residual beads were removed using the magnetic stand. 25% total volume (300 μL) trichloroacetic acid was added, vortexed, and incubated at 4°C overnight for precipitation. Next day, proteins were pelleted at maximum speed at 4°C for 30 min, washed with 1 mL of cold acetone, and air dried after removing acetone at RT for 1 min. Proteins were solubilized in 10 μL of 1xLDS buffer with 30 mM DTT and boiled at 95°C for 30 min for reverse-crosslinking. Solubilized proteins were prepared for mass spectrometry.

The protein eluate samples were taken the same volume for S-Trap (S-Trap™ micro MS sample prep kit, Protifi) procedure as protocol described. Briefly, protein solution was denatured with 10% SDS, reduced with 10 mM final concentration of DTT at 60 ℃ for 15 minutes and alkylated with 20mM final concentration of IAM at room temperature for 30 minutes. Then, all samples were quenched with 10 mM DTT to eliminate excess IAM in the samples. The protein samples were then acidified with 27.5% phosphoric acid to reach pH ≤ 1. Proteins were trapped into the S-Trap column by centrifuge at 10,000 g for 30 seconds and washed with 100 mM TEAB (final) in 90% methanol repeatedly. 1 μg Trypsin/LysC was added to protein for overnight digestion at 37 °C. Digested samples were quenched with 0.2% formic acid. Samples were then eluted from the S-Trap column with sequential addition and centrifuge of buffer 1 (50 mM TEAB), buffer 2 (0.2% formic acid) and buffer 3 (50% acetonitrile). The eluted solution was pooled and subsequently dried by SpeedVac (SavantTM SpeedVacTM SPD120, Thermo Fisher Scientific).

Each dried sample was resuspended with 80 µL 100mM HEPEs (pH 8.5) with gently vortex and centrifuge. 8 new TMT labels were resuspended with 80 µL ACN, (OptimaTM, LC/MS grade, Fisher ChemicalTM.) and 20 µL of TMT reagent was added to corresponding sample. All samples were incubated in room temperature for 1 hour. 1 µL of each TMT labelled sample was taken out and mixed to perform a label check, which served as a quality control (QC) for efficacy of TMT labeling process. The TMT label check mixture was mixed with LC buffer A (0.1% Formic Acid) and injected into the mass spec. LC/MS run of label check sample indicated the label efficiency is about 99%. TMT labelled samples were mixed and dried down with Thermo SpeedVac. (SpeedVac SPD120, Thermo Fisher Scientific). Combined samples were desalted using C18 stage tips (Cat # PTR-92-05-18, PhyNexus Inc.).

The dried peptides samples were reconstituted with LC Mobile phase A and analyzed by nano flow HPLC (Ultimate 3000, Thermo Fisher Scientific) followed by Orbitrap EclipseTM TribridTM (Thermo Fisher Scientific). Nanospray FlexTM Ion Source (Thermo Fisher Scientific) was equipped with Column Oven (PRSO-V2, Sonation) to heat up the nano column (Aurora Ultimate, 250 mm x 75 µm ID, 1.7 µm C18, IonOpticks) for peptide separation. The nano LC method was water acetonitrile based 120 minutes long with 0.3 µL/min flowrate. All TMT labeled peptides were first engaged on a trap column (Cat. No: 164535, Thermo Fisher Scientific) and then were delivered to the separation nano column by the mobile phase. A TMT specific MS2-based mass spectrometry method on Orbitrap Eclipse was used to sequence TMT peptides that were eluted from the nano column.

The ionized peptides were fractionated by FAIMS Pro™ using a 3-CV (−45, −60, −75 V) method. For the full MS, 120,000 resolution was used with the scan range of 400 m/z – 1600 m/z. ‘Standard’ AGC target and ‘Auto’ Maximum Injection Time were selected. For the dd-MS(MS2), resolution was 50,000 and isolation window was 0.7 Da. Normalized AGC Target was set at 250%. Maximum Injection Time Mode was Auto and Collision Energy mode was ‘Fixed’. TMT Quant was performed with Proteome Discoverer 2.5. Raw data was searched using UniProt Homo Sapiens database, Proteome ID UP000005640.

### Total MS

Samples were analyzed on the timsTOF Ultra (Bruker Daltonics, Germany) coupled to a nanoElute 2 (Bruker Daltonics, Germany) with 50 ng sample input. Samples were eluted off an ionoptiks Aurora Ultimate Ultra C18 column (25 cm length x 75 µm ID x 1.7 um particle size; ionoptiks, Australia) at 50 °C with an integrated 10 µm emitter inside a nanoelectrospray Captive Spray source (Bruker Daltonics, Germany) with a 60 min active gradient (2-26% Solvent B in 48-mins, 26-35% Solvent B in 12 mins, 35-95% Solvent B in 0.10 mins, wash at 95% for 7.90 mins; Solvent B: 0.1% formic acid in acetonitrile, Solvent A: 0.1% formic acid). The TIMS 1/K0 mobility range was 0.75-1.30 V·s/cm2, with a ramp time of 50 ms and 100% duty cycle. The source was set to 1600 V, 3.0 l/min dry gas, and 200°C dry temp. diaPASEF windows were created with py_DIA^109^ with a mass range of 350-1250 Da and a mobility range of 0.60-1.45 1/K0 giving an estimated cycle time of 2.25 s. The collision energy ramped from 20 to 59 eV at 0.60 to 1.60 V·s/cm2.

The .d files acquired in data-independent mode were analyzed using Spectronaut v 18.2 (Biognosys AG, Schlieren, Switzerland) with the directDIA+ feature against the Uniprot Homo sapiens protein database. Proteolysis with Trypsin was assumed to be semi-specific, allowing for N-ragged cleavage with up to two missed cleavage sites, with a PSM false discovery rate (FDR) of 1%. Data filtering was based on the Q Value, and global normalization was utilized. Cysteine modified with iodoacetamide was set as a fixed modification. Variable modifications included oxidation of methionine and tryptophan, deamidation of asparagine and glutamine, and phosphorylation of serine, threonine, and tyrosine, acetylation of N-terminal and lysine, and pyro-glutamic acid conversion from glutamine and glutamic acid.

### AlphaFold3-based structural analysis

A modeling technique which combined *de novo* structure prediction from AlphaFold 3^67^ and manual refinement^68^ was used to develop a three-dimensional structure of the protein-RNA complex of SRSF1 and the exon 1 mRNA of *PVT1*. To begin the structural modeling, the canonical protein sequences of SRSF1 (UniProt: Q07955) was used. Specifically, to constrain the challenge of *de novo* structure prediction and to improve the accuracy, the protein sequence was reduced to only those sub-sequences or domains which were annotated to be interacting with RNA. Thus, for SRSF1 the residues 10–200 were selected as they contained the RNA recognition motifs (RRM), i.e., RRM1 and RRM2 connected by an inter-domain linker. Similarly, the 5’ 80 nt sequence from *PVT1* exon 1 was used as input. The seed used for the prediction was 1234567 and only the default five predicted models were used for further screening. The models were visualized and screened using UCSF ChimeraX^110^, of which a model that had the best concordance with known annotation was chosen as the representative structure.

### DeepCLIP score analysis

SRSF1 binding score was calculated by using a deep learning method, DeepCLIP^69^. This method was trained on eCLIP SRSF1 data from ENCODE-K562 cell line after which the query wildtype and mutant sequences of *PVT1* exon 1, each of length 75nt were provided. The mutant sequences contained a GGA > UUA substitution.

### eCLIP analysis

The processed eCLIP bigWig tracks of SRSF1 and SRSF7 performed in the K562 and HepG2 cell lines were downloaded from the ENCODE database. eCLIP tracks were visualized using Integrative Genomics Viewer (v2.19.1).

### MYC rescue assay

Conditional MYC expressing murine hepatocellular carcinoma cells, EC4 cells were seeded at a cell density of 7000 cells per well in 96 well plate. Cells were transfected with the plasmid constructs (200ng/well) after 24 h (Day 1). Following transfection, MYC transcription was turned off treating cells with doxycycline (2ng/mL) after 48 h of transfection (Day 3). The cell viability was measured using CellTiter-Glo Luminescent Cell Viability Assay (Promega, G7570) to evaluate the rescue effect of the plasmid constructs. The luminescence values were normalized to GFP construct and the relative change in cell viability was calculated. The statistical significance was calculated using ordinary one-way ANOVA in GraphPad Prism 10.2.2.

### CRISPRi PVT1 RNA-seq

Lentivirus was generated using the UCOE-SFFV-dCas9-BFP-KRAB construct (Addgene #85969). COLO320DM cells were transduced with the lentivirus, incubated for 2 days, and sorted for BFP-positive cells via flow cytometry. Polyclonal BFP-positive cells were subsequently monocloned to isolate clones with stable and homogenous expression of dCas9-BFP-KRAB and high CRISPR interference (CRISPRi) efficiency. The guide RNAs for PVT1 (sgPVT1) and non-targeting control (sgNTC) were used as described in Hung et al. Nature 2021^4^.

COLO320DM dCas9-KRAB BFP cells were plated 24 hours prior to transduction. The cells were transduced with either sgPVT1 or sgNTC. On Day 1, 24 hours post-transduction, puromycin was added to the culture medium at a concentration of 2 µg/mL to initiate the selection process. On Day 3, 72 hours post-transduction, the cells were collected, washed in PBS, and analyzed using an Attune flow cytometer to quantify the expression and cellular effects of the sgRNA targeting. On Day 4, 96 hours post-transduction, the cells were harvested, and RNA was extracted for bulk RNA sequencing. RNA libraries were constructed using TruSeq Stranded Total RNA Library Prep Kit with Ribo-Zero (Illumina, catalog no. 20020596) and sequenced by Nextseq 550 (Illumina), 75 bp paired-end reads per sample.

### CRISPRi PVT1 RNA-seq analysis

Raw paired-end RNA-seq fastq files were aligned to the GRCh38 genome reference using STAR (version 2.7.10b). Following alignment, read counts for each gene were obtained using featureCounts (version 2.0.8) for subsequent analyses. From these read counts, differentially expressed genes between CRISPRi sgPVT1 and sgNTC (non-targeting control) COLO320DM cells were identified using the DESeq2 R package (version 1.44.0). The gene list, ordered by log2 fold change as calculated by DESeq2, was then used for Gene Set Enrichment Analysis (GSEA) with the GSEA function from the clusterProfiler R package (version 4.12.2), employing human hallmark gene sets from the msigdbr R package. Significantly enriched pathways were defined as those with adjusted p-values less than 0.05, adjusted using the Benjamini-Hochberg method.

Expression levels of canonical *MYC* and *PVT1-MYC* were assessed by FPKM (Fragments Per Kilobase of transcript per Million read). Reads that were aligned to MYC exon 1 or those spanning *MYC* exon 1–2 were classified to canonical *MYC* transcripts while reads spanning *PVT1* and *MYC* exon 2–3 were classified to *PVT1-MYC*. Assigned read counts were normalized by the length of canonical *MYC* and *PVT1-MYC* transcripts, respectively and total read counts per million. Following this, we normalized RNA expression levels of canonical *MYC* and *PVT1-MYC* FPKM in CRISPRi sgPVT1 divided by corresponding sgNTC to evaluate relative expression changes.

### Quantification and statistical analysis

Details of exact statistical analysis, tests, and other information can be found in the main text, figure legends, and Methods.

## Supplemental information (4)

**Table S1. Fusion transcript information, related to Figure 1**

Information of fusion transcripts detected in TCGA+CCLE databases and cell line long-read RNA-seq. 1) RNA fusion transcripts from the cancer samples with matched WGS and RNA-seq in TCGA and CCLE databases (n=1,825) with amplicon type annotation and expression level (FFPM). 2) Fusion transcripts detected by long-read RNA-seq in ecDNA(+) cell lines and their isogenic cell lines with breakpoint and spanning read counts.

**Table S2. Lists of oligo sequences used in this study, related to Figures 2–4**

List of oligo sequences used in this study. 1) RT-qPCR primers. 2) ChIRP probes. 3′ Biotin-TEG is denoted as O at the end of each oligo sequence. 3) Flex-seq custom probes. LHS (Left-hand side) and RHS (Right-hand side) probe sequences are shown along with the combined full-length sequence of probe.

**Table S3. Total MYC ChIRP-MS and total MS, related to Figure 3**

Identified proteins in total MYC ChIRP-MS and total MS. 1) Total MYC ChIRP-MS in COLO320DM and COLO320HSR cells. Normalized protein abundance by RNase control ChIRP and z-score of each protein with gene name are shown. 2) Normalized protein abundance across cell lines (COLO320DM, COLO320HSR, HCT116, and IMR90) of each protein with gene name are shown (NaN; no detection).

**Table S4. Flex scRNA-seq and CRISPRi PVT1 RNA-seq, related to Figure 4**

Differentially expressed genes in 1) Flex scRNA-seq in COLO320DM cells (*in vitro*) and 2) COLO320DM xenograft (*in vivo*) model. Differentially expressed genes were identified comparing the Q5 (highest) vs Q1 (lowest) quintiles of *PVT1-MYC* expression. 3) Differentially expressed genes in COLO320DM CRISPRi *PVT1* RNA-seq. Differentially expressed genes were identified comparing the gene expression in the sgPVT1 and sgNTC samples.

